# Genetic basis and evolution of structural color polymorphism in an Australian songbird

**DOI:** 10.1101/2023.09.03.556140

**Authors:** Simon Yung Wa Sin, Fushi Ke, Guoling Chen, Pei-Yu Huang, Erik Enbody, Jordan Karubian, Michael S. Webster, Scott V. Edwards

## Abstract

Island organisms often evolve phenotypes divergent from their mainland counterparts, providing a useful system for studying adaption under differential selection. Some island birds have melanic plumage differing from the color of mainland conspecifics, a trait proposed as an insular adaptation. In the white-winged fairywren (*Malurus leucopterus*), subspecies on two islands have a black nuptial plumage whereas the subspecies on the Australian mainland has a blue nuptial plumage. The black subspecies have a feather nanostructure that could produce a blue structural color, suggesting a blue ancestor. An earlier study proposed independent evolution of melanism on the islands based on the history of subspecies divergence. However, the genetic basis of melanism and the origin of color differentiation in this group are still unknown. Here, we used whole-genomes to investigate the genetic basis of melanism by comparing the blue and black *M. leucopterus* subspecies to identify highly divergent genomic regions. We identified a well-known pigmentation gene *ASIP* and four candidate genes that may contribute to feather nanostructure development. We also detected signatures of a selective sweep in genomic regions containing *ASIP* and *SCUBE2* not in the black subspecies, as predicted by earlier work, but in the blue subspecies, which possesses many derived SNPs in these regions, suggesting that the mainland subspecies has re-evolved a blue plumage from a black ancestor. This re-evolution was likely driven by a pre-existing female preference. Our findings provide new insight into the evolution of plumage coloration in island versus continental populations, and, importantly, we identify candidate genes that likely play roles in the development and evolution of feather structural coloration.

## Introduction

Studies of the genetic basis underlying phenotypic variation are crucial to understanding processes leading to evolutionary adaptation. The “island syndrome”, which describes the divergence of organisms on islands from their continental counterparts^1^, has provided model systems ideal for studying adaptive phenotypes under differential selective pressures^2,3^. For example, insular populations of birds frequently have evolved plumage coloration differing from related continental populations^4,5^. One well-known example is that of island melanisms, where populations of birds^4,6–8^ and other taxa^9–11^ on islands evolve melanic coloration that is speculated to confer an advantage^12–17^. The repeated evolution of melanic coloration in island populations provides an opportunity to understand the genetic basis of adaptive evolutionary convergence, yet knowledge of the underlying molecular mechanisms of this convergence remains limited to a few case studies^4,6,8^.

An intriguing case of island melanism occurs in the white-winged fairywren (*Malurus leucopterus*), which has intra-specific variation in plumage structural color rather than pigment-based coloration^7,18^. The color polymorphism in this species could originate due to loss or gain of structural coloration^7,19,20^. Although we have made good progress in identifying genes responsible for pigment-based plumage coloration in birds in recent years, such as pigmentation due to melanins^21^, carotenoids^22^, and psittacofulvins^23^, the genetic basis of feather structural colors is still largely unknown^24^. Structural colors are produced by coherent scattering or interference of light by nanostructures with periodic variation in refractive index^25^. Many birds have iridescent (changing in hue depends on the viewing or illumination angle) or non-iridescent structurally based plumage colors^26,27^. Non-iridescent structural plumage colours are produced by coherent light scattering due to quasi-ordered β-keratin nanostructures and air in the medullary spongy layer of feather barbs^28^, which generates blue, violet, and ultraviolet feather colors. The presence of melanosomes (melanin-containing organelles) beneath the spongy layer is also essential for the production of structural color as it absorbs incoherently scattered light^27^.

*Malurus* fairywrens (Maluridae) are passerines displaying considerable variation in ornamental structural colors of males, from blue, indigo, violet, to almost pure UV hues^29^. Most species exhibit a high degree of sexual dichromatism, and breeding males have striking ornamentation, whereas females of most populations are cryptically brown in coloration. Structural color variation in fairywrens is associated with the thickness of the keratin cortex on top of the spongy layer, the size and density of the spongy layer, the relative amount of melanosomes underneath the spongy layer, and their interactions^29^. This variation in different microstructural components contributing to the diversity of structural colours in fairywren nuptial plumages makes them an ideal model system to study mechanisms of structural colour evolution.

The white-winged fairywren (*Malurus leucopterus*) is particularly well-suited to studying the genetic basis of structural coloration because males in this species have either blue or black nuptial plumage. *Malurus leucopterus* is distributed across most of Australia, with males of the mainland subspecies (*M. l. leuconotus*) having a bright cobalt blue nuptial plumage with white wings, whereas the two island subspecies in Western Australia (*M. l. leucopterus* on Dirk Hartog Island [DHI] and *M. l. edouardi* on Barrow Island [BI]) exhibit a black male nuptial plumage with white wings^7,18^ (Fig. 1). DHI and BI are far (∼600 km) from each other and separated from continental Australia by ∼2 km and ∼56 km, respectively. The feather barbs of blue males on the mainland are composed of a spongy layer with a keratin matrix and air space, with melanin granules beneath the spongy layer around the central vacuole: a feather nanostructure typically producing blue colors^18,30^. Surprisingly, feather barbs of black males on both DHI and BI also have a spongy layer that could in principle produce a blue color similar to that on the mainland^18,30^, but the feather barbs of these males have several features that render them black instead of blue in coloration: they have a thicker cortex with more melanin; a spongy layer with more and larger holes; and melanin granules that are distributed both within and beneath the spongy layer^18,30^. Because black feather barbs in other species do not normally possess a spongy layer, the presence of a spongy layer that could produce blue color in black feathers might suggest that the black *M. leucopterus* island populations evolved from blue ancestors^18,30^. The low divergence between *M. leucopterus* populations (*F*_ST_ = 0.04 - 0.18)^31^ indicates that the switch of male nuptial plumage color occurred quickly and in the recent past, implying a simple genetic change^7^.

**Figure 1.**
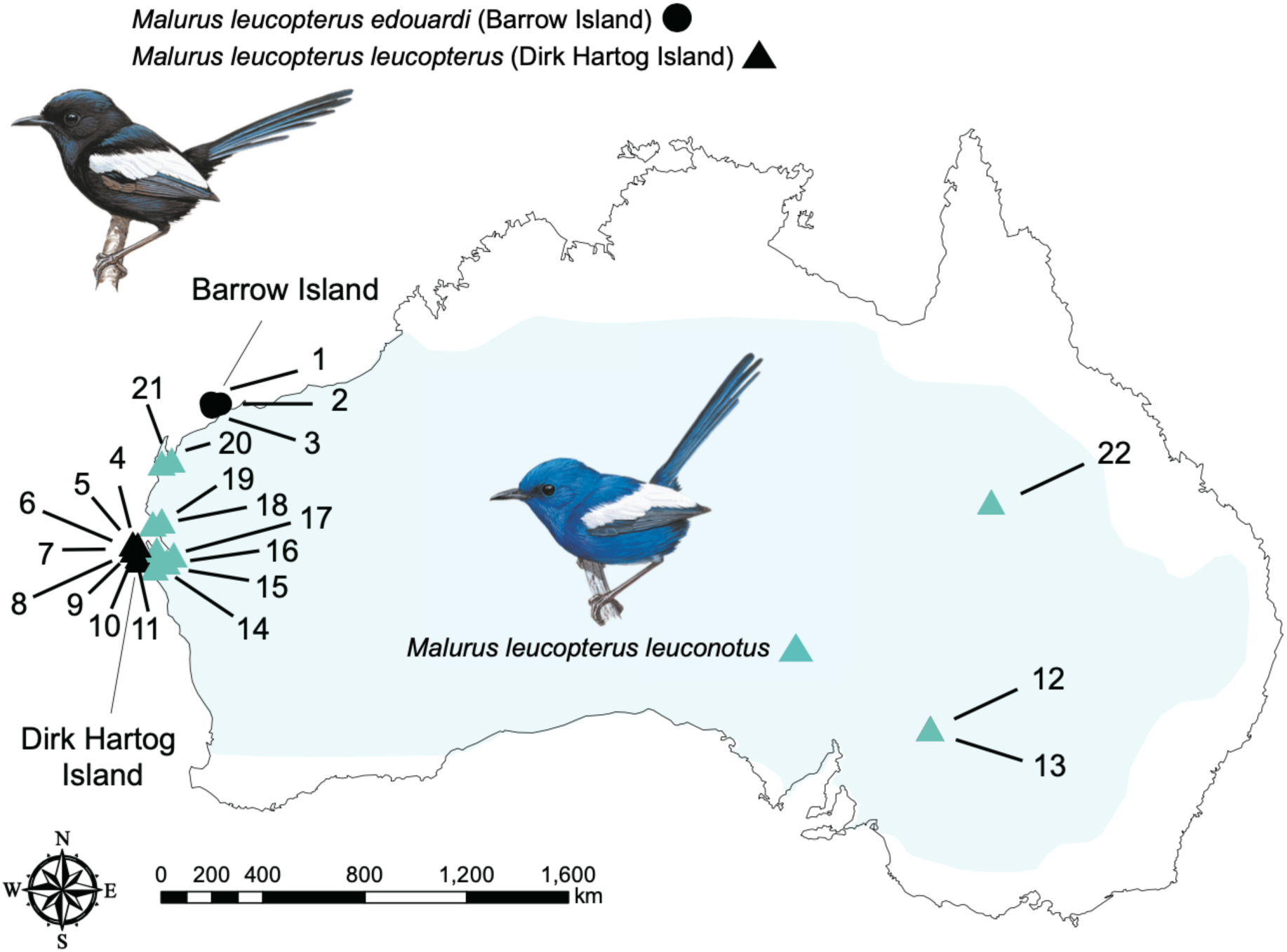
Geographical distribution and sampling sites of *Malurus leucopterus*. *M. leucopterus* males from Barrow Island (*M. l. edouardi*, denoted by black circles on the map) and Dirk Hartog Island (*M. l. leucopterus*, denoted by black triangles) have black nuptial plumage. *M. leucopterus* males distributed across the Australian continent (*M. l. leuconotus*, denoted by blue triangles) are with blue plumage. Light blue color indicates the range of *M. leucopterus*. Refer to Supplementary Table S1 for detailed information of sampled individuals. Illustrations of birds were reproduced with the permission of Lynx Edicions.

*Malurus leucopterus* subspecies on DHI and mainland have been suggested based on reduced representation sequencing data to be more closely related to each other than either is to the subspecies on BI^31^. The emergence of color polymorphism in *M. leucopterus* therefore occurred either due to convergent melanism on the two islands or because the mainland subspecies evolved blue plumage from a black ancestor (Fig. 2)^7,19,20^. The two sister species in the bi-colored wren clade, *M. melanocephalus* and *M. alboscapulatus*, found on mainland Australia and New Guinea, respectively, also have black male nuptial plumage, which could imply a black ancestor to the bi-colored wren clade, but feather microstructure suggests this plumage pattern arose independent of the switch(es) to melanism in *M. leucopterus*^30^. Based on the independent colonization events of DHI and BI, and the presence of spongy layer in black subspecies suggesting a blue ancestor, Walsh et al.^31^ concluded that there was independent evolution of melanic plumage in island *M. leucopterus* subspecies. However, the genetic basis of melanism, both for *M. leucopterus* and for other species in the bi-colored clade, is still an unknown. Accordingly, the alternative hypothesis is still plausible, such that melanism on the islands could occur with or without convergence, even if colonization occurred independently (Fig. 2), and understanding the genetic basis of black plumage in this clade will help clarify mechanisms of plumage color evolution generally^32^.

**Figure 2.**
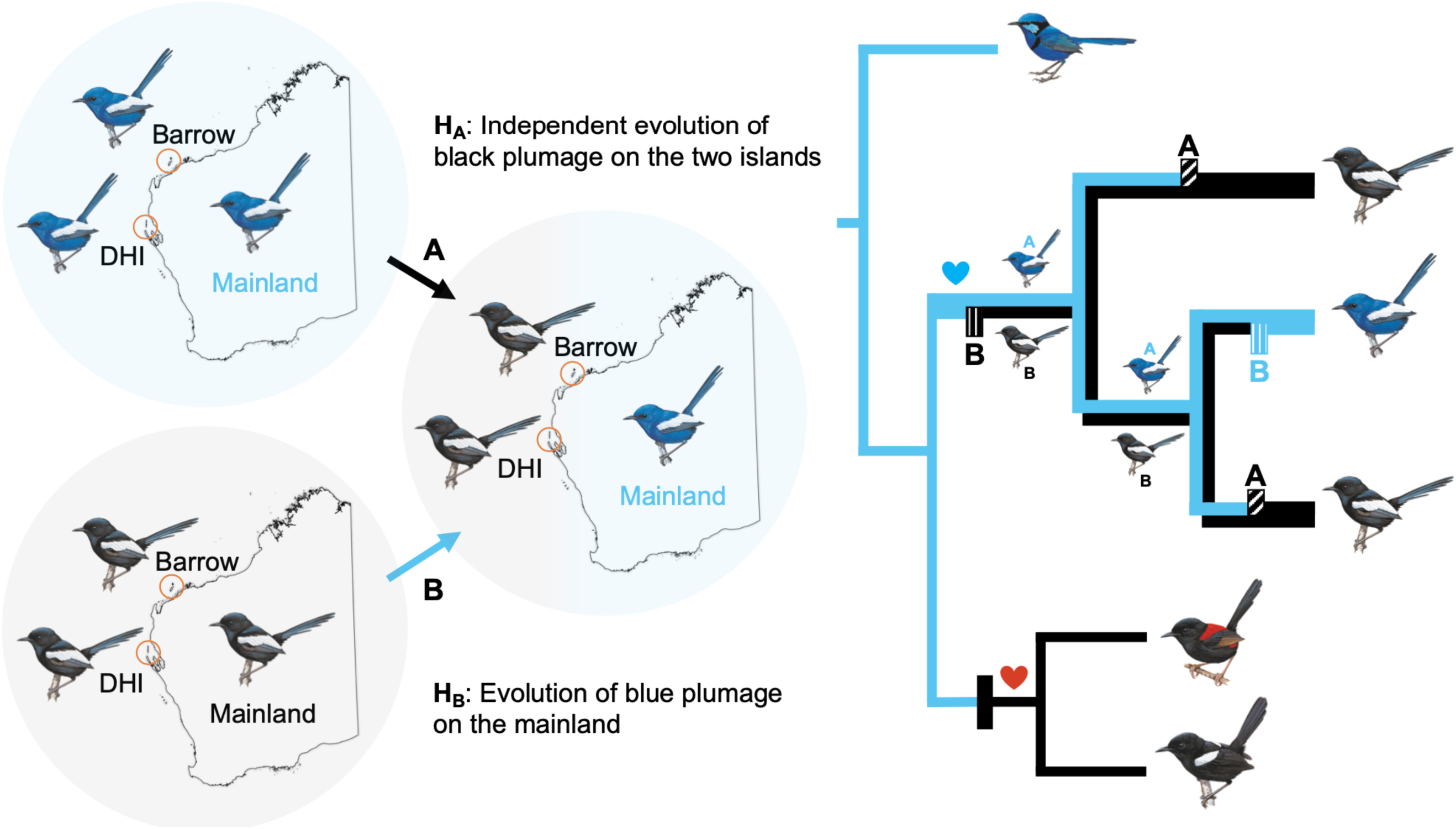
Hypotheses of plumage color evolution in the white-winged fairywren (*Malurus leucopterus*). Left side of the figure shows the two hypotheses: hypothesis A (H_A_) – independent evolution of black plumage on Dirk Hartog Island (DHI) and Barrow Island (BI); hypothesis B (H_B_) – evolution of blue plumage on the mainland from birds with black plumage. Right side shows the corresponding positions of the plumage color change on the phylogenetic tree predicted by the two hypotheses. In the *M. leucopterus* clade, the upper half of the branches leading to the subspecies indicates scenario under hypothesis A, and the lower half of the branches represents hypothesis B. Hypothesized color changes are indicated by boxes (H_A_: with diagonal lines and “A” above, H_B_: with vertical lines and “B” below, black: change from blue to black, blue: change from black to blue). The heart symbol denotes proposed female color preference based on male petal carrying behavior. Female preference for blue color likely maintained in all *M. leucopterus* subspecies, and female preference for red color might have evolved in the common ancestor of *M. melanocephalus* and *M. alboscapulatus*. Illustrations of birds were reproduced with the permission of Lynx Edicions.

Here we used whole-genome sequencing to study the genetic basis of melanism in *M. leucopterus*. We compared the blue and black *M. leucopterus* subspecies to identify highly divergent genomic regions between subspecies and candidate genes associated with plumage color differentiation. We further looked for signatures of selective sweeps at these loci and also examined genotypes in *M. leucopterus* subspecies and other *Malurus* species to test hypotheses of plumage color evolution in *M. leucopterus*.

## Results

### Phylogeny of the bi-colored clade and *M. leucopterus* subspecies

Phylogenomic analysis identified a monophyletic clade of *M. leucopterus* (Fig. 3), sister to the clade formed by two other bi-colored wrens, *M. melanocephalus* and *M. alboscapulatus*. Within *M. leucopterus*, the two black-plumaged insular populations did not form a monophyletic clade (Fig. 3). The maximum-likelihood (ML) tree (Fig. 3A) shows that *M. l. edouardi* from BI formed a monophyletic clade that was sister to the intermixed cluster of individuals from the mainland (*M. l. leuconotus*) and DHI (*M. l. leucopterus*). In this cluster, *M. l. leucopterus* formed a monophyletic group, while individuals with blue plumage from the mainland were paraphyletic (Fig. 3A). The coalescent-based SNAPP tree (Fig. 3B) instead shows that *M. l. edouardi* with black plumage was more closely related to blue-plumaged *M. l. leuconotus* than to black-plumaged *M. l. leucopterus*. Both trees therefore indicate that the two insular populations were divergent from each other (Fig. 3 and S1), and one of the black-plumaged insular populations was more closely related to the blue-plumaged mainland population than to another black-plumaged population.

**Figure 3.**
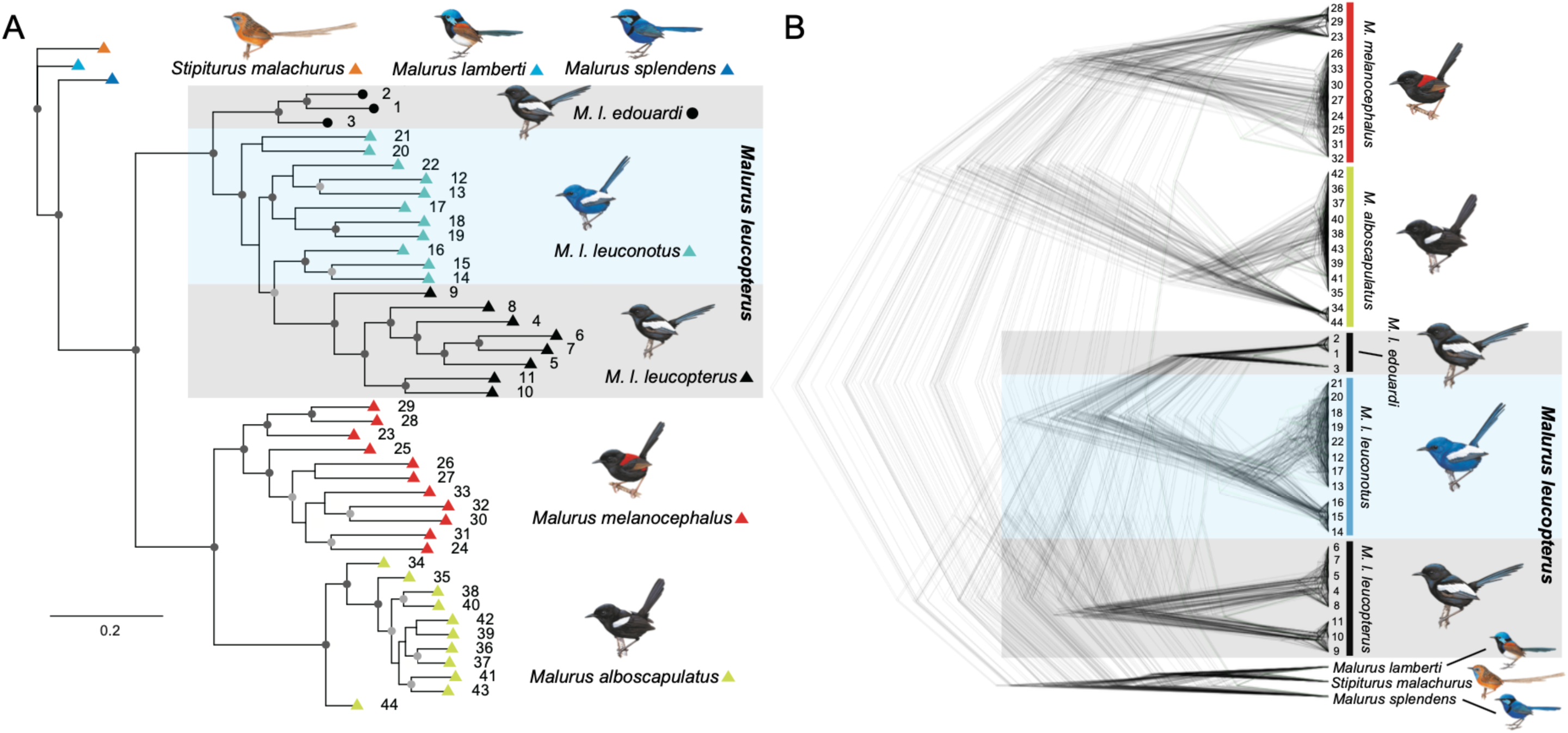
Phylogenetic positions of *Malurus leucopterus* subspecies. (A) Maximum-likelihood phylogenetic tree based on 168,378 nuclear SNPs. Branches with dark grey dots indicate bootstrap values of 100, and light grey dots are with bootstrap values >80 and <100. (B) SNAPP tree based on 6,790 nuclear SNPs. Illustrations of birds were reproduced with the permission of Lynx Edicions.

### Genomic regions of divergence reveal possible targets of selection for male plumage color

Genome-wide divergence (mean autosomal *F*_ST_) between *M. l. leuconotus* (blue plumage) and *M. l. leucopterus* (black plumage) was 0.119, and between *M. l. leuconotus* and *M. l. edouardi* (black plumage) was 0.274 (Fig. S2). In addition to comparison between subspecies, combining the two subspecies with black plumage (i.e. *M. l. leucopterus* and *M. l. edouardi*) in the divergence analysis facilitated identification of divergent genomic regions between blue and black birds, lowering the divergence (mean autosomal *F*_ST_) between black insular and blue mainland subspecies to 0.081 (Fig. 4). Using windowed *F*_ST_ estimates, we identified 42 highly divergent genomic regions (exceeding 99.9^th^ percentile) relative to the background (Fig. 4; Table S2), of which 20 were autosomal. These divergent genomic regions contained 90 annotated genes (Table S2) enriched for gene ontology categories of osteoblast and muscle cell differentiation and migration (Fig. S3) and not showing any obvious link to plumage color development. However, five genes – *ASIP*, *SCUBE2*, *THBS2*, *THBS4*, and *NECTIN3* – were plausibly associated with plumage color differentiation (Fig. 5). *ASIP* has a known regulatory role in vertebrate melanogenesis. *SCUBE2*, *THBS2*, *THBS4*, and *NECTIN3* all play roles in keratinocyte development and cell-to-cell/matrix interactions. When we separated the two black subspecies for the *F*_ST_ analysis, the comparison between *M. l. leuconotus* and *M. l. leucopterus* shows that the same regions holding these five genes were all above the 99.9^th^ percentile (Fig. S2B), whereas only the regions containing *ASIP* (scaffold 285) and *SCUBE2* (scaffold 132) passed the cut-off in the comparison between *M. l. leuconotus* and *M. l. edouardi* (Fig. S2C).

**Figure 4.**
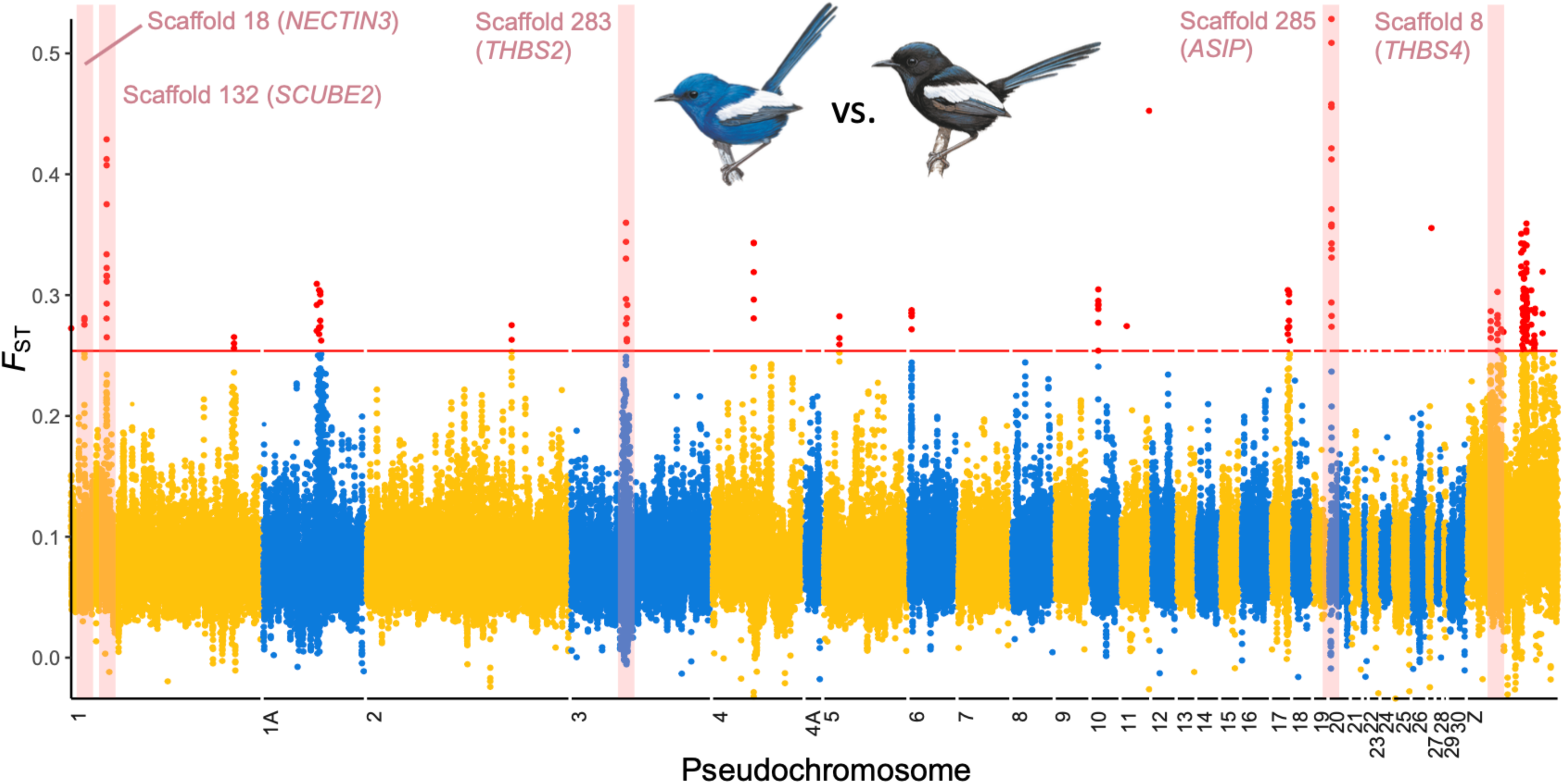
Genome-wide differentiation between black and blue *Malurus leucopterus*. Points indicate overlapping sliding window *F*_ST_ values in 50 kb windows. Red points above red horizontal line are above 99.9^th^ percentile. Five divergent regions with genes that are potentially relevant to plumage color development are highlighted in pink, with scaffold and gene identities labeled. Scaffolds are reordered as pseodochromosomes according to the *Taeniopygia guttata* genome. Illustrations of birds were reproduced with the permission of Lynx Edicions.

**Figure 5.**
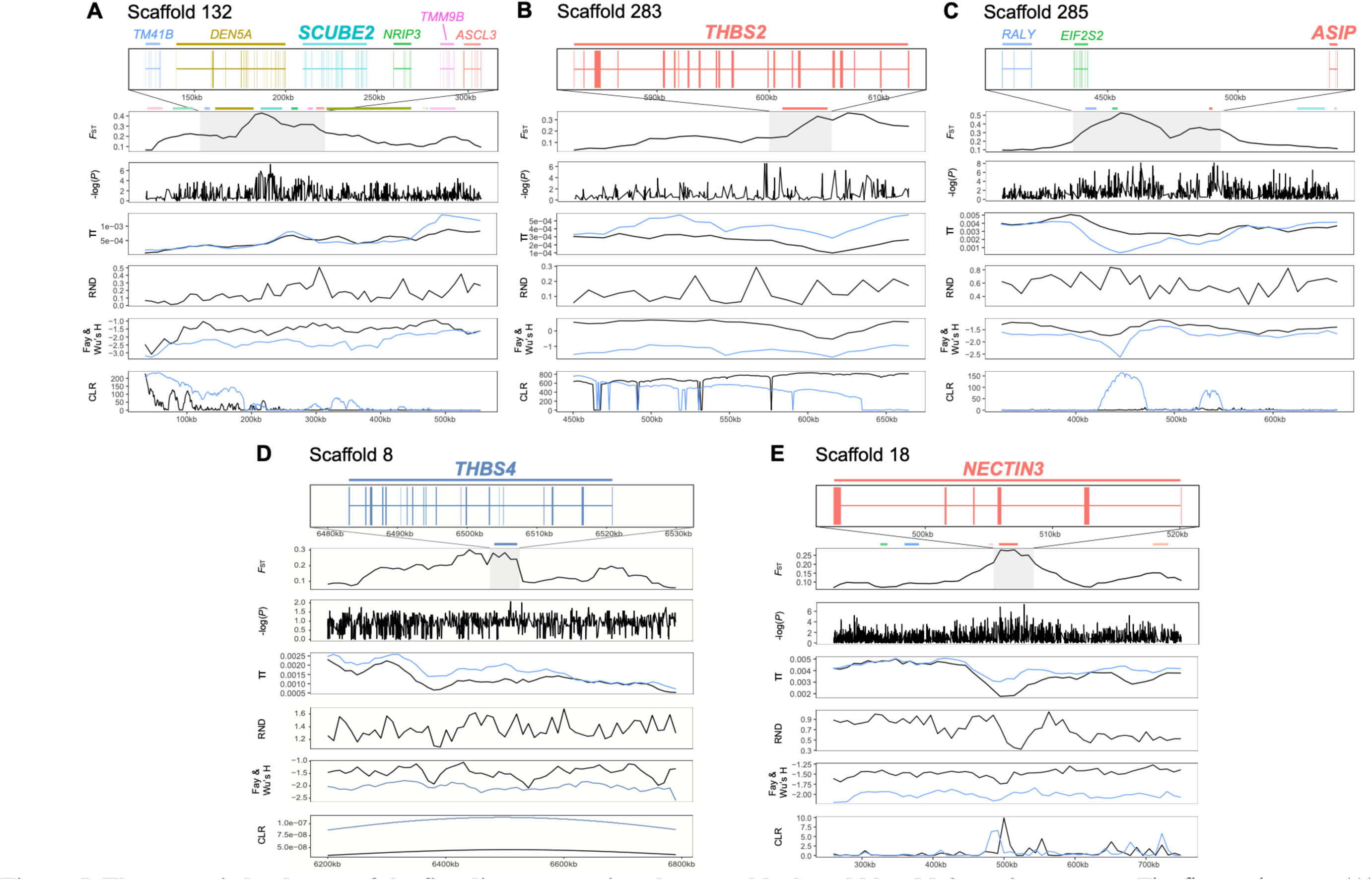
The genomic landscape of the five divergent regions between black and blue *Malurus leucopterus*. The five regions are (A) scaffold 132 containing *SCUBE2*, (B) scaffold 283 containing *THBS2*, (C) scaffold 285 containing *ASIP*, (D) scaffold 8 containing *THBS4*, and (E) scaffold 18 containing *NECTIN3*. *F*_ST_, SNP associations, genetic diversity (π), relative node depth (RND), Fay and Wu’s H, and composite likelihood ratio (CLR) are shown in different panels for each region. Blue and black lines in the same panel of π, Fay & Wu’s H, and CLR represent blue and black populations, respectively. Annotated genes in those regions are depicted using blocks and lines to represent exons and introns, respectively, in the top panel.

In the five highly divergent regions, SNPs strongly associated with plumage color were identified by GWAS in the loci and/or in the regions flanking those loci (Fig. 5). The region with the *ASIP* gene was lower in nucleotide diversity in the blue (π < 0.001; Fig. 5) than black birds (π > 0.002), which was in turn lower than the global π (autosomal; blue subspecies = 0.0034; black subspecies = 0.0033). No introgression was identified from blue wrens to the blue *M. leucopterus* in those regions based on the low relative node depth (RND) values (Fig. 5). Signatures of selective sweep were detected in the *ASIP* and *SCUBE2* regions in the blue birds, indicated by the significant negative Fay and Wu’s H values (<99th percentile^33^, i.e. −2.59 of blue population) in blue birds and higher SweepFinder2 composite likelihood ratio (CLR) values in blue than black birds (Fig. 5). Strikingly, the phylogeny of these two genomic regions differed from the species tree (Shimodaira-Hasegawa test, p < 0.001; Fig. 6A and 6B, Fig. S4): in these trees individuals clustered by plumage color. This could be due to the selective sweep in the blue birds. The genotype matrix of those two regions also show a pattern of selective sweep in the blue birds (Fig. 6C and 6D), which was not detected in the birds with black plumage.

**Figure 6.**
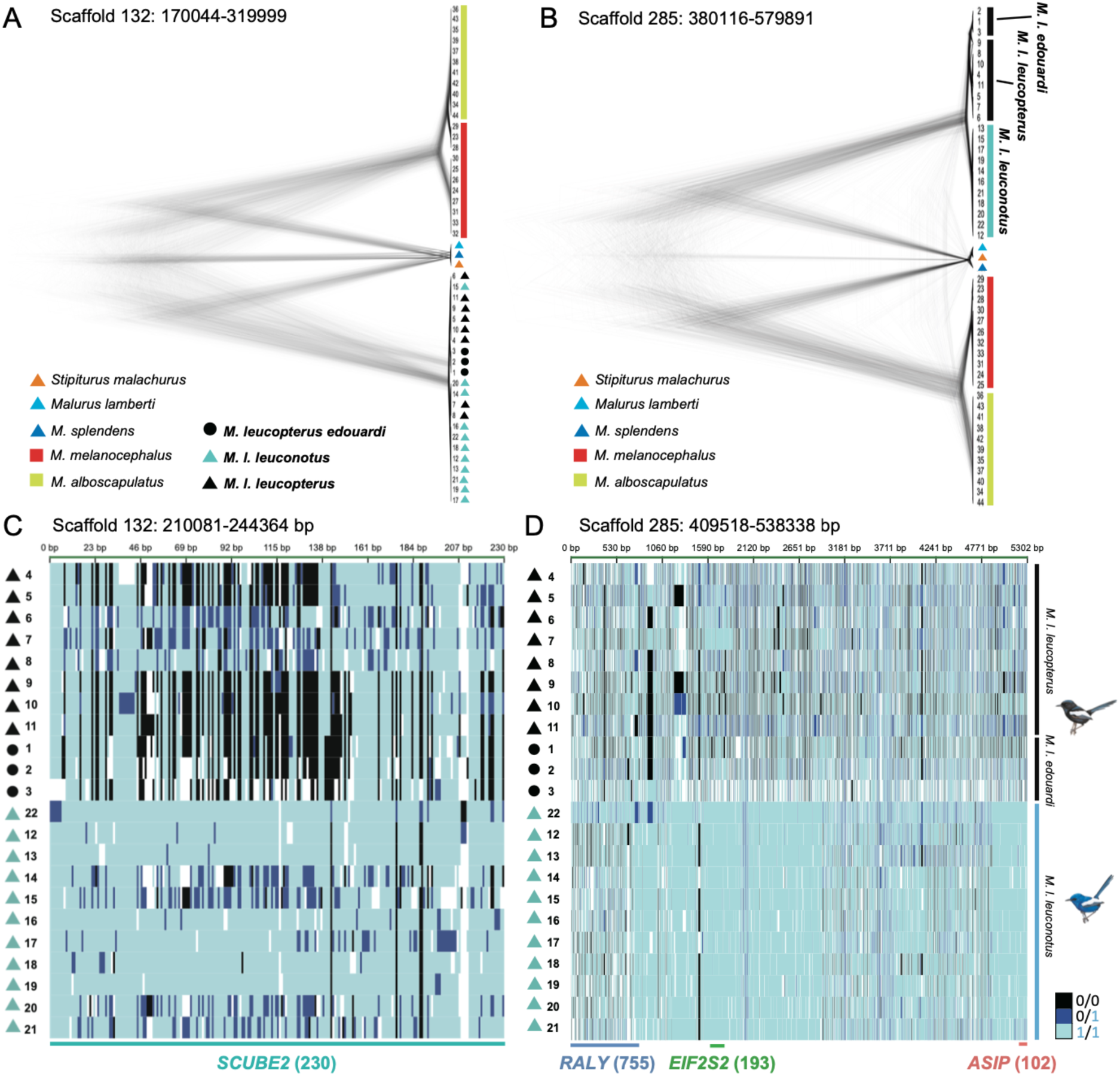
Phylogeny and individual genotypes of the two regions of divergence. Phylogenetic tree reconstructed using SNAPP based on SNPs from the whole divergence peak region contains (A) *SCUBE2* or (B) *ASIP*. (C-D) Genotypes at SNPs between black and blue *Malurus leucopterus* populations in the (C) *SCUBE2* and (D) *ASIP* regions. Each row represents one individual. Black triangles and circles denote *M. leucopterus* from Dirk Hartog Island and Barrow Island, respectively. Light blue, black, and dark blue indicate positions homozygous for the allele that was the same as the reference genome (from a *M. leucopterus leuconotus* sampled in Queensland, Australia), homozygous for the allele different from the reference, and heterozygous for both alleles, respectively. Missing values are represented by white. Number in parentheses indicates the polymorphic site number of each gene. Illustrations of birds were reproduced with the permission of Lynx Edicions.

When we extracted those loci that were highly associated with plumage color differentiation between DHI and mainland populations in the five highly divergent genomic regions and compared the genotypes and allele frequencies between BI and DHI, the pattern for scaffold 285, 283, and 132 (1 out of 2 SNPs) was similar between the two insular populations (Fig. 7). The genotypes and allele frequencies of the mainland population were different in these regions, especially for the region in scaffold 285, which was consistent with the selective sweep signature in the blue birds. The comparison of genotypes among all sequenced bi-colored wrens and outgroup individuals showed that the selective sweep regions of the blue *M. leucopterus* (i.e. scaffold 285 and 132) contain many derived SNPs that are different from the other two bi-colored wrens and outgroups (Fig. 7). Moreover, the SNPs in black *M. leucopterus* were not found in the other two bi-colored wrens that were also black in male nuptial plumage color, suggesting that the evolution of black plumage in *M. leucopterus* was independent from that in *M. melanocephalus* and *M. alboscapulatus*.

**Figure 7.**
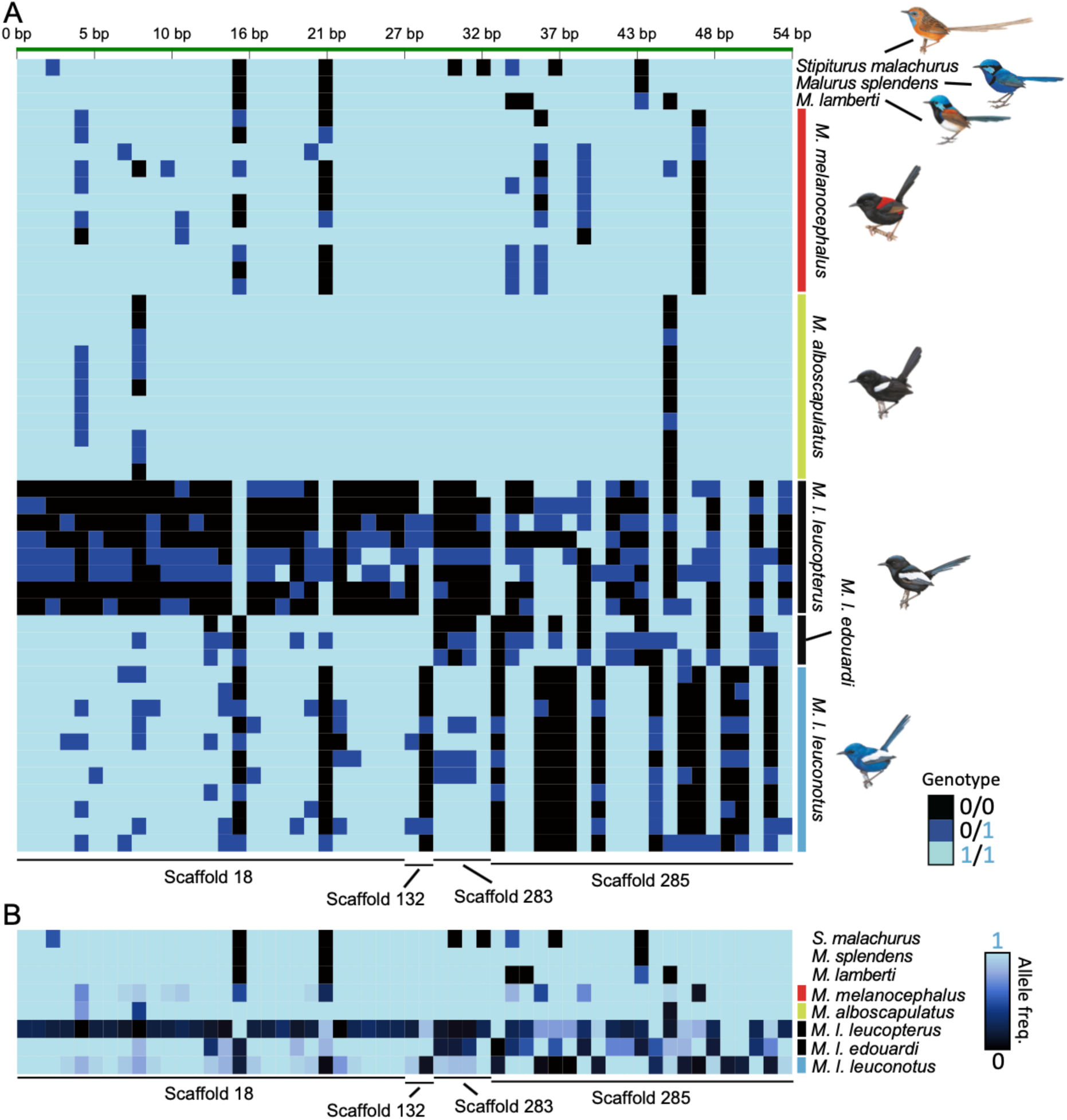
(A) Genotypes and (B) allele frequencies at 54 SNPs in the five highly divergent genomic regions (see Fig. 5) that were strongly associated (-log(*P*)>6) with plumage color differentiation between *Malurus leucopterus leucopterus* from Dirk Hartog Island and *M. leucopterus leuconotus* from the mainland. (A) Genotypes of *M. leucopterus edouardi* from Barrow Island, other bi-colored wrens, and outgroups are also shown. Each row represents one individual. Light blue, black, and dark blue indicate positions homozygous for the allele that was the same as the reference genome (from a *M. leucopterus leuconotus* sampled in Queensland, Australia), homozygous for the allele different from the reference, and heterozygous for both alleles, respectively. Each column represents one SNP position. The scaffolds where the SNPs originated from are indicated at the bottom. (B) Each row represents the allele frequencies of one species or subspecies of *M. leucopterus*, ranging from 0 (all alleles were different from the reference) to 1 (all alleles were the same as the reference). Illustrations of birds were reproduced with the permission of Lynx Edicions.

## Discussion

### Genetic basis of plumage color difference in *M. leucopterus*

We have identified highly divergent genomic regions between blue and black *M. leucopterus* subspecies containing candidate genes associated with plumage color development (Fig. 4 and 5). The candidate genes include an important pigmentation gene and genes that may play a role in keratinocyte and feather nanostructure development. Variation of structural color in fairywrens is related to multiple independent features of the feather barb rather than a single factor^29^. Both the density and distribution of melanin and thickness and nanostructure of cortex and spongy layer contribute to chromatic and achromatic variation in structural color^29^. The gene encoding the ASIP (agouti signaling protein), which is an antagonistic ligand of MC1R, is in the most divergent genomic region between blue and black *M. leucopterus* and is the only pigmentation gene identified in the divergent regions. ASIP is an important melanin production modifier that can switch the synthesis of eumelanin to pheomelanin and regulate melanocyte precursor behavior in hair follicles^34^.

In birds, *ASIP* was found to be up-regulated in peripheral pulp adjacent to apigmented feather barbs, and it suppresses melanocyte differentiation from progenitor cells during feather growth^35^. Differential melanocytes are required to produce and transport melanosomes to keratinocytes in developing feathers^27,35,36^, an expression of *ASIP* would therefore inhibit the formation and deposition of melanosomes. *ASIP* expression has been shown to affect body color and pattern in birds and other vertebrates^37–43^. Mutation of this gene has also been reported to link to island melanism in *Monarcha castaneiventris*, which has an insular population possessing a non-synonymous substitution in the CDS of *ASIP*^6^. In contrast, no mutations in the *ASIP* CDS were fixed in *M. leucopterus* subspecies (supplementary results). The upstream region of *ASIP* is highly divergent between blue and black subspecies and likely regulates the expression of this gene. Mutations in the genomic region containing *ASIP*, *EIF2S2*, and *RALY* are also associated with pigmentation change in other avian species^37,41^. We predict *ASIP* to be underexpressed in developing feathers of black *M. leucopterus* relative to the blue form, which leads to more differential melanocytes in developing feathers and hence more melanosomes in both the cortex and spongy layer of the feather barbs, resulting in the black color of the feather^35^.

Another candidate gene that is also in a divergent genomic region with a selective sweep signature and may link to structural plumage color development is *SCUBE2* (signal peptide, cubulin [CUB] domain, epidermal growth factor [EGF]-like protein 2). This gene can interact with Sonic Hedgehog (SHH) and enhance SHH signaling activity^44^, and it plays important roles in SHH-related biology such as organ development^45^, including feather development^46^. SHH induces feather development and mediates programmed cell death in the feather barb marginal plate epithelium during keratinization of the barb plate epithelium^47,48^, which regulates barb and barbule formation^49,50^. *SCUBE2* may play a role in developing feather barb nanostructure required for structural color production, and accounting for the difference in the nanostructure between blue and black subspecies (e.g. a more degraded spongy layer in black feathers). The peak of the genomic island of divergence is at the upstream region of *SCUBE2*, suggesting mutations in a cis-regulatory region may affect gene expression. While protein difference could also be linked to the differentiation in phenotype, since a nearly fixed non-synonymous substitution between blue and black subspecies has also been identified (supplementary results).

Other candidate genes that are in divergent genomic regions and may link to feather development and plumage color difference are *NECTIN3*, thrombospondin-2 (*THBS2*), and thrombospondin-4 (*THBS4*). Nectin-3 is a cell-cell adhesion molecule and it plays an important role in the formation of cell-cell adhesion junctions in keratinocytes^51^. Nectin-3 is required for cell adhesion and differentiation in hair follicles and for proper hair formation in mice, and *NECTIN3* mutants have impaired cell adhesion and abnormal hair^52^. Thrombospondins are extracellular glycoproteins participate in cell-cell and cell-extracellular matrix interaction^53,54^. Both *THBS* genes are important for skin keratinocyte development^53–55^. These three genes are only divergent between DHI and mainland populations but not between BI and mainland populations (Fig. S2), which could be due to the small sample size from BI or independent evolution of these genes on the two islands. With their associations with keratinocyte development, these genes, plus *SCUBE2*, may play a role in developing the feather nanostructure required for structural color, which is an important step in unravelling the molecular mechanism and evolution of structural coloration in birds.

### Re-evolution of blue plumage in mainland *M. leucopterus*

We identified genomic islands of divergence that show a signature of selective sweep (Fig. 5 and 6). Surprisingly, selective sweeps occurred in the genomic regions containing *ASIP* and *SCUBE2* in the blue mainland subspecies but not the black subspecies on islands, in contrast to signatures of selective sweep found in insular *M. castaneiventris* populations showing convergent evolution of island melanism^6^. Individual genotypes and allele frequencies in these two sweep regions in the blue subspecies are different from the two island subspecies (Fig. 7), which are more similar to each other even though they are more divergent genome-widely. Given the similarity of the two black subspecies in these two regions, the ancestor of *M. leucopterus* subspecies could have already become melanic before they arrived DHI and BI, i.e. black plumage was ancestral for all the subspecies. The two island subspecies possess derived SNPs in these two regions that are not found in *M. melanocephalus* and *M. alboscapulatus*, which also have evolved black male nuptial plumage. If these are, or are closely linked to, the causal mutations render the plumage black in color in the two *M. leucopterus* island subspecies, the switch to melanism in the other two bi-colored wrens is an independent event. This independent origin of melanism is supported by the different extents of feather nanostructure degradation – *M. melanocephalus* has a high degree of structural degradation of the spongy layer in the black feather barb, whereas black *M. leucopterus* subspecies possess a spongy layer that is similar in nanostructure to those of blue *M. leucopterus* and *M. splendens*^30^, suggesting the switch to melanic form happened later in *M. leucopterus* than the other two melanic bi-colored species.

The degradation of feather nanostructure in these black fairywrens is consistent with an initial hypermelanization, followed by subsequent changes in spongy layer keratin organization and ramus shape for pigment presentation^30^. The proposed hypermelanization happened in the common ancestor of all *M. leucopterus* subspecies could be due to mutations in the *ASIP* region, which were likely inherited by all subspecies. This explains the sharing of SNPs in that region by the two island populations but no other species. Hypermelanization would then have relaxed selection to maintain the feather nanostructure needed for structural color, hence genes contributing to the nanostructure might accumulate mutations that affect the phenotype, which was shown by the spongy layer degraded with more holes ^30^. If the melanic form was at the same time favored by natural selection, feather nanostructure would be selected for features that consolidate the black color, such as the thicker cortex and rami observed^30^. These changes could be due to mutations in *SCUBE2* or other candidate genes relevant to feather nanostructure development, and could occur before or after the divergence of the subspecies. However, different from the other two black bi-colored wren species, black *M. leucopterus* still have a functional spongy layer that can produce a blue color by coherent light scattering^18^, which means only a reversal of hypermelanization was required to switch the black plumage to blue again. The strong selective sweep signature in the genomic region containing *ASIP* supports this hypothesis of re-evolution of blue plumage on the mainland by mutations that reversed hypermelanization, in contrast to the hypothesis of convergent melanism on islands^31^. This sweep region in the blue subspecies contains many derived SNPs associated with the plumage color difference between blue and black subspecies (Fig. 7). These derived SNPs are not only different from the two island subspecies but also from the other two bi-colored wrens and outgroups, including one of the blue wrens *M. splendens*. The mainland subspecies likely re-evolved the mechanism to develop blue plumage again, from a black *M. leucopterus* ancestor that evolved from a blue ancestor. Assuming *SCUBE2* is link to the development of feather nanostructure, the selective sweep happened in the genomic region containing this gene in the blue subspecies was likely to postdate the de-hypermelanization, since selection could only act on to favour the development of optimal feather nanostructure for blue structural color after the density and distribution of melanosome were altered. The mutations of the *SCUBE2* region might facilitate the development of feather nanostructure similar to its blue ancestor or the sister blue wren species, e.g. a more structured and intact spongy layer^30^, in order to produce a more coherent light scattering and brighter blue hue^29^.

Therefore, instead of a case of convergent evolution of melanism on island populations^31^, the differentiation in plumage color between the island and mainland populations could be a case of re-evolution of blue plumage in the mainland population. The mutations and selective sweep in these two genomic regions containing important candidate genes, especially the pigmentation gene *ASIP*, in *M. l. leuconotus* was likely the mechanism of re-evolution of blue plumage on the mainland. Although hybridizations between fairywren species have been reported^56–58^, including between *M. l. leuconotus* and the Superb fairywren *M. cyaneus*^58^, re-evolution of blue plumage in mainland *M. leucopterus* was not a result of genetic introgression from blue wrens (Fig. 5). The high diversity in feather plumage color among *Malurus* species and body regions within a species suggest that plumage color is a very labile trait during evolution of these birds^29^, and re-evolution of blue plumage from a black one is feasible. A substantial difference in plumage color can be due to minor changes in the size and structure of light-scattering elements or pigment concentration or distribution^25,29,59^. Convergent regulatory changes in melanin synthesis pathway have also led to repeated evolution of structural coloration in birds^27^. In *Malurus*, Fan et al.^29^ showed that black feather barbs from species that have both black and blue plumage patches, such as *M. splendens* and *M. lamberti*, contain a spongy layer that could produce a blue color, but the degradation and high density of melanosome in the spongy layer plus the thinker and more melanized cortex make them appear black. This suggests that evolutionary changes in modular regulatory control of color for blue and black plumage patches to be common and relatively feasible in this clade, which likely also applies to the switches between blue and black whole body plumages in *M. leucopterus*. The mainland population of *M. leucopterus* diverged from the DHI population in late Pleistocene^31^, when the population size began to expand on the mainland (supplementary results; Fig. S5). This rapid population expansion might increase dispersal and gene flow on mainland and facilitate the spread of mutations for a “novel” phenotype, i.e. the blue plumage, especially if the trait was under strong selection.

### Selective pressure for plumage color differentiation between islands and mainland

*Malurus* fairywrens are one of the birds with the highest rates of extra-pair paternity^60–62^, and their high sexual dichromatism and very colorful male plumages indicate that their plumage is under strong sexual selection^63,64^. Males with brighter plumage are more attractive to females and have higher within- and extra-pair paternity^60,65^. In *M. leucopterus*, female preference for bright blue color is likely retained from their ancestor in all descendant subspecies. Males of many fairywren species often carry colorful petals in their bills during courtship towards females^66,67^. The black male *M. leucopterus* on the islands have higher incidence of petal carrying than mainland males and show a strong preference to carry blue petals^16^, suggesting female’s preference for males carrying blue petals. Female’s preference for bright blue color has likely been maintained from their ancestor that had a blue male nuptial plumage, even though it became black (Fig. 2), and could be the selective force driving the re-evolution of blue male nuptial plumage in the mainland *M. leucopterus* population. If males carrying the mutations for a bluer plumage have a higher reproductive success, then the haplotype would spread quickly in the population, leaving a signature of selective sweep^68^ in *M. l. leuconotus*. We therefore propose that the pre-existing female preference for blue color in *M. leucopterus* had driven the re-evolution of blue nuptial plumage.

In contrast to the mainland subspecies, black plumage on the two islands is maintained, even though there is gene flow between mainland and island subspecies^31^. The maintenance of black plumage on islands could be due to weaker sexual selection on islands and/or natural selection. Both sexual dichromatism and dimorphism of body size in this species are reduced on islands^16,69^. Small population size, low genetic diversity, and limited choice of mates are believed to reduce sexual selection in isolated island populations compared to mainland counterparts^16,70–72^. The effective population size on the islands was estimated to be much smaller than that on the mainland^31^. In this study, the autosomal π were 0.0030 and 0.0019 for the DHI and BI subspecies, respectively, which were lower than the 0.0034 of the mainland subspecies. In addition to the difference in sexual dichromatism, mainland birds have bigger clutches, shorter incubation periods, and higher realised reproductive success^16^. Island birds also have fewer helpers and are largely socially monogamous, and importantly appear to have lower levels of extra-pair paternity^16^. These differences in reproductive behaviour and social structure could further result in reduced strength of sexual selection on islands compared to mainland. In parallel, black plumage may confer an advantage under natural selection through providing solar radiation protection^12^, bacterial degradation resistance^13^, abrasion resistance^14^, thermoregulation^15^, and crypsis against predation and brood parasitism^16,17^. However, since black plumage might have evolved on the mainland but not small islands, similar to what likely happened in the other two bi-colored wrens, natural selection specific to insular environments alone cannot explain the evolution of black plumage. Alternatively, indirect selection through pleiotropic effects was proposed to be the evolutionary driver of melanism^31^. It is worth noting that the estimated size of the ancestral *M. leucopterus* population before splitting into three subspecies is much smaller than the current mainland population and comparable to the current DHI population^31^, suggesting that the strength of sexual selection could be lower in the ancestral population than the later expanded mainland population and might have led to a reduced sexual dichromatism in the ancestral form.

## Conclusions

Feather barb nanostructure and melanosome features are crucial component in avian structural coloration. Their nature of being independent and evolutionary labile are likely to facilitate the evolution of a high diversity of plumage structural color, both within and between species^29,73^. Feather structural colors have independently evolved many times in more than 45 avian families^27^. The convergent evolution of similar feather barb nanostructure and melanosome distribution in many avian lineages shows that the genetic basis to develop the medullary spongy layer and change melanosome density and distribution in keratinocytes is simple, and feather structural color can evolve relatively easily^27^. The transition to blue structural color, from the black plumage *M. leucopterus* that already possess a partly degraded spongy layer interrupted by melanin granules, appears to be an example of this evolutionary lability. We show that a selective sweep in the genomic region containing the pigmentation gene *ASIP* was likely the cause of a change in the melanism state of the feather barb, which led to the re-evolution of the blue plumage color. Any changes in the molecular mechanisms that affect melanin synthesis and deposition would influence the potential to develop plumage structural color. In *M. l. leuconotus*, genetic changes related to feather nanostructure development likely happened following the mutations in the *ASIP* region to consolidate the blue structural color. The parallel evolution of non-iridescent structural color in many avian lineages^27^ could follow the same order of evolutionary changes. Future studies of the genetic basis in different species and how much it is shared among birds with similar feather structural color will increase our understanding of the evolution of avian phenotypic diversity and its interaction with sexual and natural selection.

## Supplemental information

Supplemental Information includes 6 figures and 3 tables and can be found with this article online.

## Acknowledgements

This research was supported by the University of Hong Kong start-up granted to S.Y.W.S. We thank Museum of Comparative Zoology, University of Washington Burke Museum, University of Kansas Biodiversity Institute, and American Museum of Natural History for providing the samples; and Tim Sackton, Leonardo Campagna, and Michaël Nicolaï for helpful discussion. The computations were performed using research computing facilities offered by the Information Technology Services at the University of Hong Kong, and the Odyssey cluster supported by the FAS Division of Science, Research Computing Group at Harvard University.

## Author contributions

Conceptualization, S.Y.W.S and S.V.E.; sample collection, J.K., E.E., M.S.W., and S.V.E.; methodology, S.Y.W.S.; analysis, S.Y.W.S., F.K., G.C., and P.Y.H.; writing – original draft, S.Y.W.S.; writing – review & editing, all authors; supervision, S.Y.W.S. and S.V.E.

## Supplemental information

### Supplementary results

#### Demographic history of *M. leucopterus*

The PSMC analysis (Fig. S5) suggested a demographic history of *M. leucopterus* characterized by an increase in *N*_e_ from ∼500,000 around 800,000 years ago (ya) to ∼1,650,000 at the beginning of Last Glacial Period (LGP; ∼110,000 ya), followed by a sharp decline. The Barrow Island population diverged right before the beginning of Pleistocene at ∼3.52 ya^31^ when the *N*_e_ of *M. leucopterus* was at a low level, which was maintained stably during early and mid-Pleistocene. The DHI population split from the mainland population at ∼346,000 ya, when the *N*_e_ started to expand. *N*_e_ of *M. leucopterus leuconotus* kept expanding after the divergence between the mainland and DHI populations and reached a maximum when LGP began, and then dropped rapidly.

#### Nearly fixed derived substitution in a gene of the blue *M. leucopterus*

We examined the CDS of the five candidate genes, and revealed one of them contains nearly fixed substitutions. Five nearly fixed nucleotide substitutions between black and blue subspecies were found in the CDS of the *SCUBE2* gene (nucleotide position 617, 1046, 1047, 1770, and 1870, Fig. S6), which included four missense mutations leading to amino acid substitutions. One of these nearly fixed substitutions involved amino acids with different properties, and the blue subspecies has the derived substitutions (Met) compared to black subspecies and other species (Thr; Fig. S6). This amino acid substitution at position 349 locates in one of the epidermal growth factor (EGF)-like domains and a cysteine-rich region. Most (7/10) blue subspecies were homozygous for the derived state and most (8/10) black subspecies were homozygous for the ancestral state (Fig. S6), while some were heterozygous, including two individuals from DHI and two individuals from Western Australia, which were from Denham and Bullara, respectively. Denham is on the mainland right opposite of DHI and Bullara is along the coast between DHI and BI. Black-plumaged *M. leucopterus* was occasionally reported on the mainland opposite DHI^74^. An individual from Denham was homozygous for the ancestral state, while no individual on island was homozygous for the derived state.

**Table S3.** De novo assembly metrics for Malurus leucopterus leuconotus and M. splendens genomes.

| Metric | *Malurus leucopterus* | *Malurus splendens* |
| --- | --- | --- |
|  | Value | Value |
| Estimated genome size | 1.02 Gb | 1.06 Gb |
| %GC content | 41.9 | 42.3 |
| Total depth of coverage | 64.6 | 60.1 |
| Total contig length (bp) | 978039748 | 978629979 |
| Total scaffold length (bp, gapped) | 1019547066 | 1003289036 |
| Number of contigs | 58256 | 47017 |
| Contig N50 | 40.8 kb | 53.4 kb |
| Number of scaffolds | 5908 | 5169 |
| Scaffold N50 | 4.29 Mb | 3.07 Mb |
| Scaffold N50 (with gaps) | 4.02 Mb | 3.08 Mb |
| Total BUSCOs | 2857/2882 (99.1%) | 2862/2882 (99.3%) |
| Complete single-copy BUSCOs | 2811/2882 (97.5%) | 2819/2882 (97.8%) |

**Figure S1.**
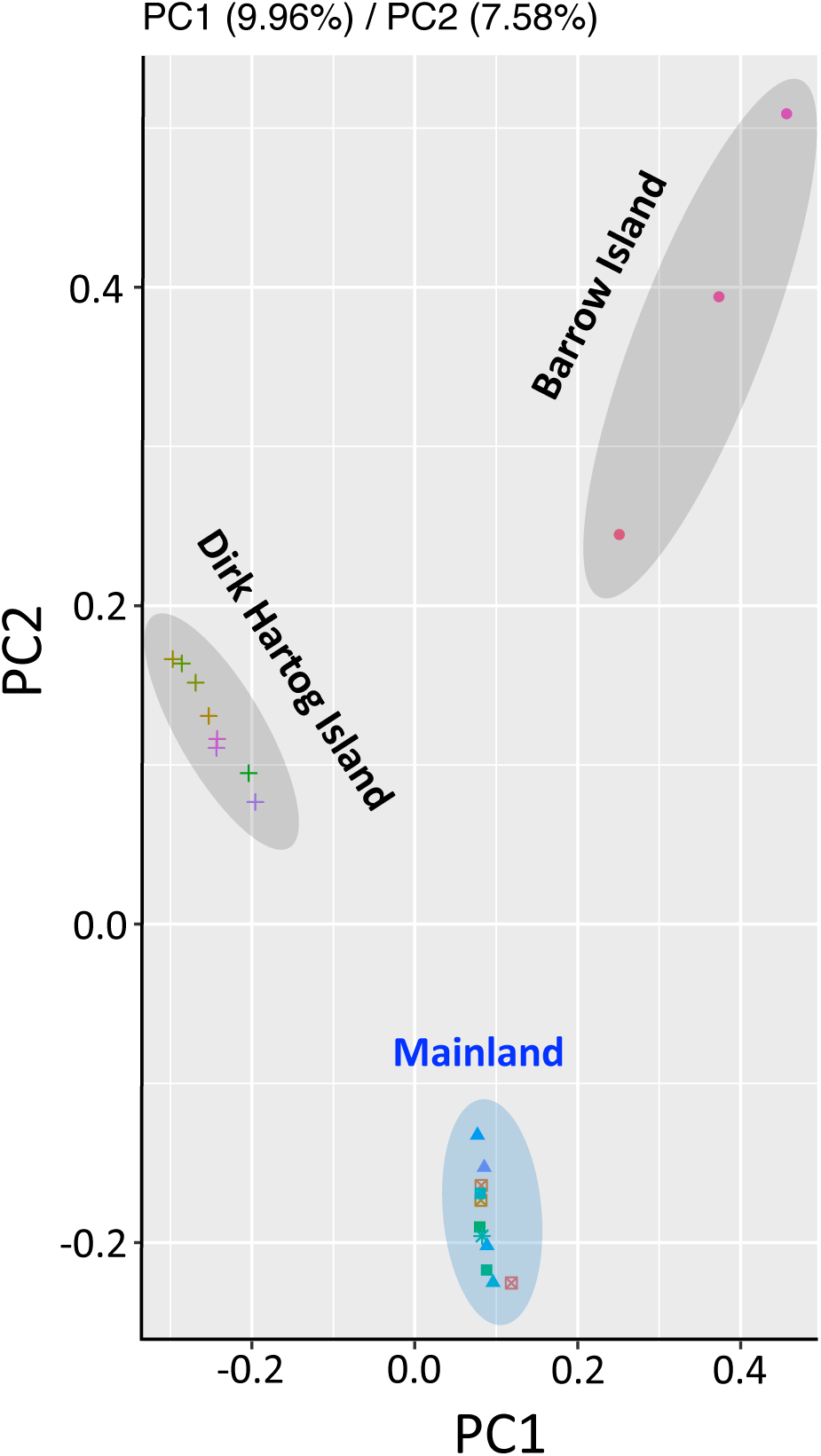
PCA of the covariance matrix generated from genotype likelihoods of all *Malurus leucopterus*. Each symbol represent one individual. The ovals denote the origin of the individual.

**Figure S2.**
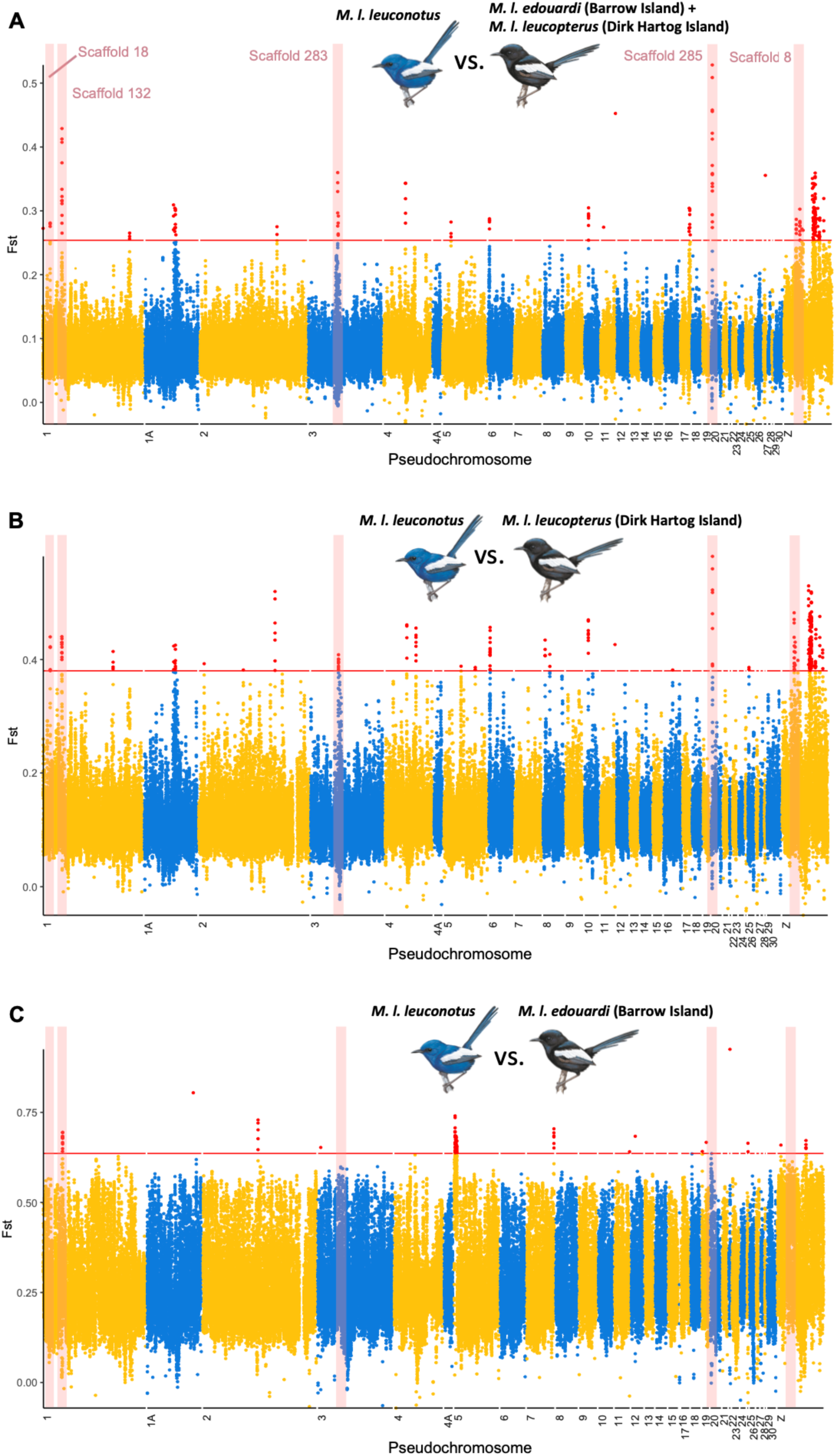
Genome-wide differentiation between *Malurus leucopterus* populations: (A) *M. l. leuconotus* and *M. l. edouardi*+*M. l. leucopterus*; (B) *M. l. leuconotus* and *M. l. leucopterus*; and (C) *M. l. leuconotus* and *M. l. edouardi.* Points indicate overlapping sliding window *F*_ST_ values in 50 kb windows. Red points above red horizontal line are above 99.9^th^ percentile. Five divergent regions with genes that are potentially relevant to plumage color development are highlighted in pink, with scaffold identities labeled. Scaffolds are reordered as pseodochromosomes according to the *Taeniopygia guttata* genome.

**Figure S3.**
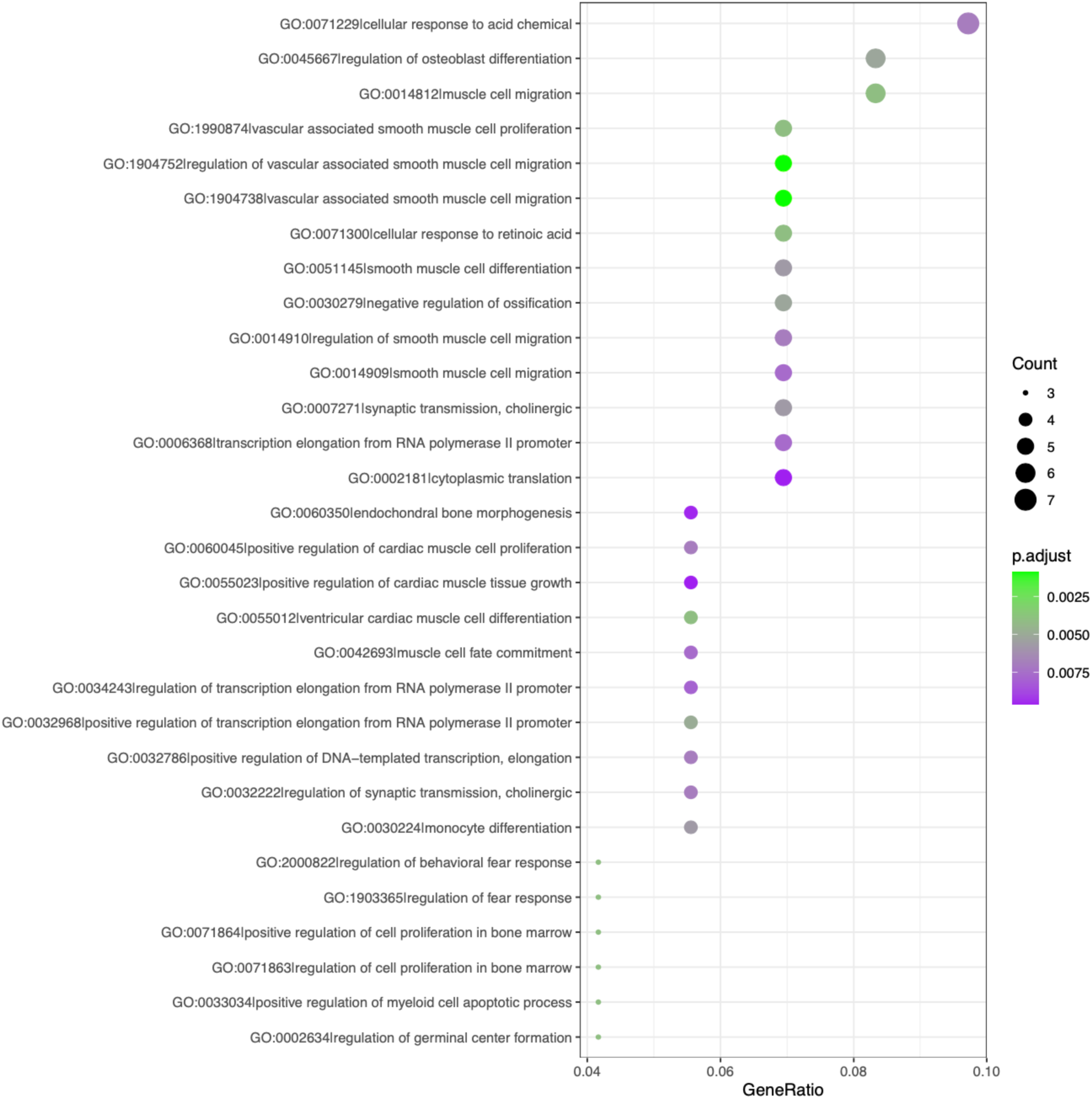
Gene ontology categories that were significantly enriched for the genes located in the top 99.9^th^ percentile *F*_ST_ peaks in the blue and black *Malurus leucopterus* subspecies comparison.

**Figure S4.**
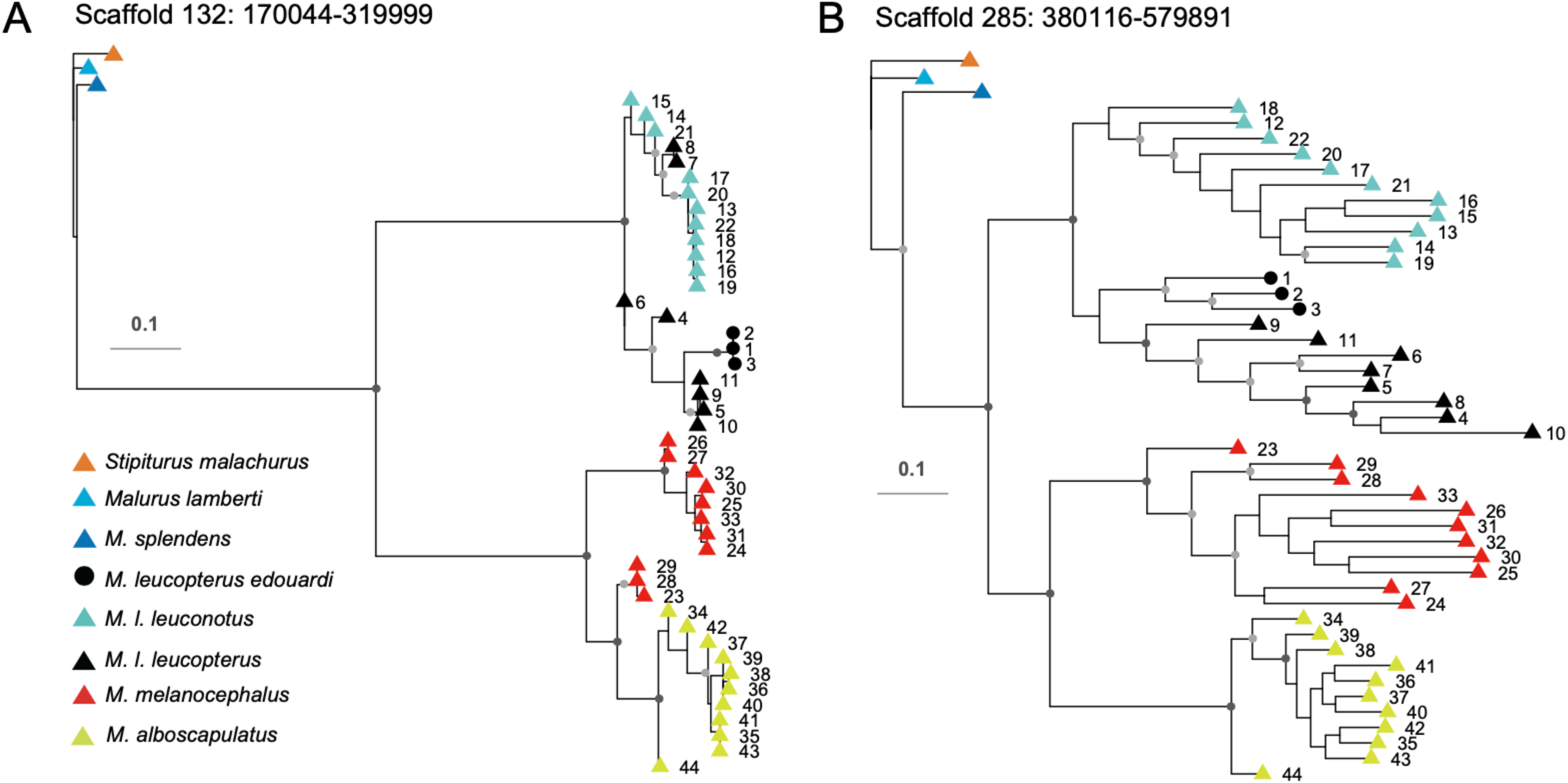
Phylogeny of the two regions of divergence. Maximum-likelihood phylogenetic trees reconstructed using SNPs from the whole divergence peak region contains (A) *SCUBE2* or (B) *ASIP*. Branches with dark grey dots indicate bootstrap values of 100, and light grey dots indicate bootstrap values >80 and <100.

**Figure S5.**
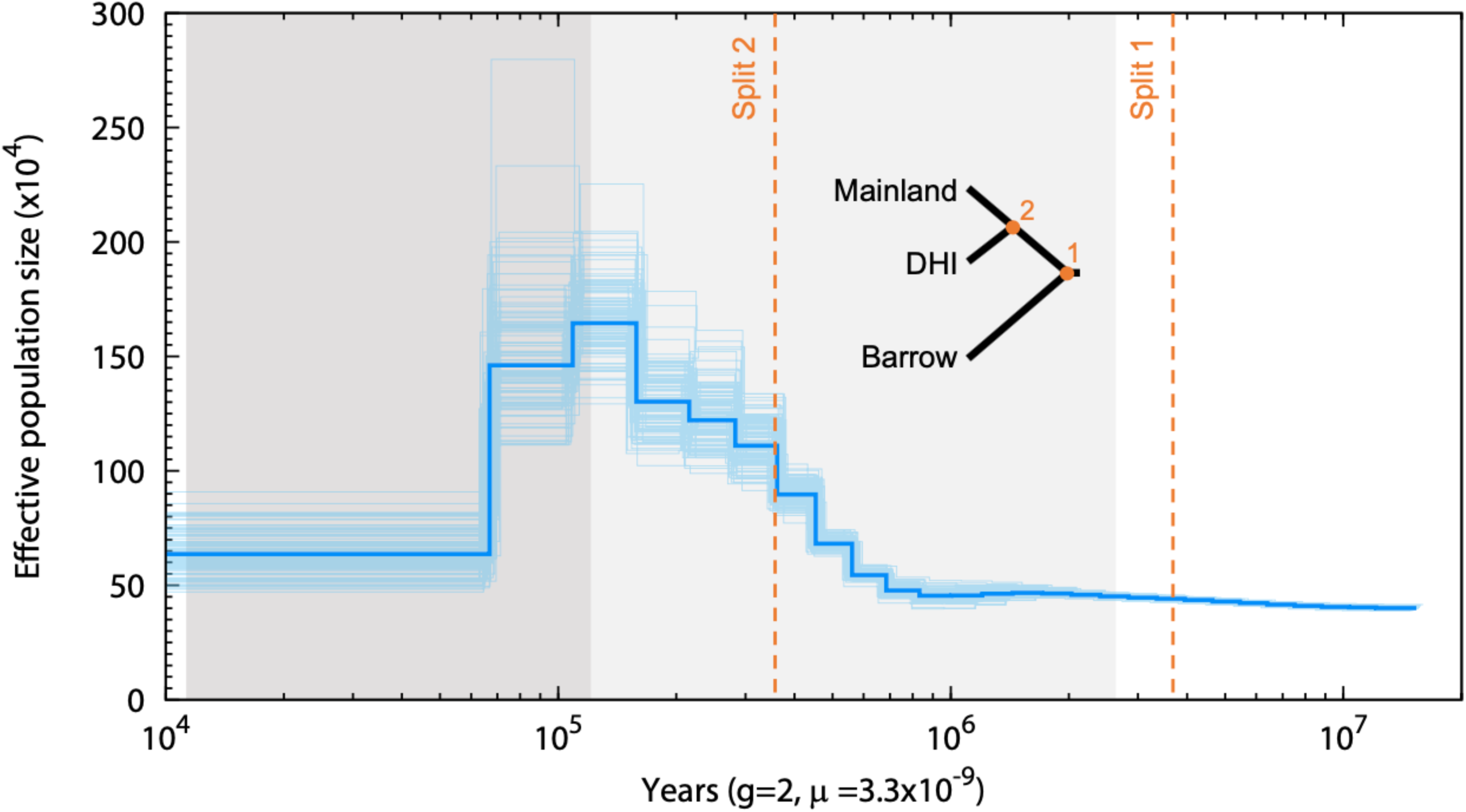
Demographic history of the white-winged fairywren (*Malurus leucopterus*) inferred using PSMC. Bold blue line is the median effective population size estimate, whereas thin lines are 100 individual bootstrap replicates. The topology shows the phylogenetic relationships of the three subspecies. The orange dotted lines indicate the estimated time of the divergence between the mainland subspecies and Dirk Hartog Island subspecies (Split 2) and the split between their common ancestor and Barrow Island subspecies (Split 1) (Walsh, et al. 2021). The light and dark gray areas indicate the Pleistocene epoch and Last Glacial Period, respectively.

**Figure S6.**
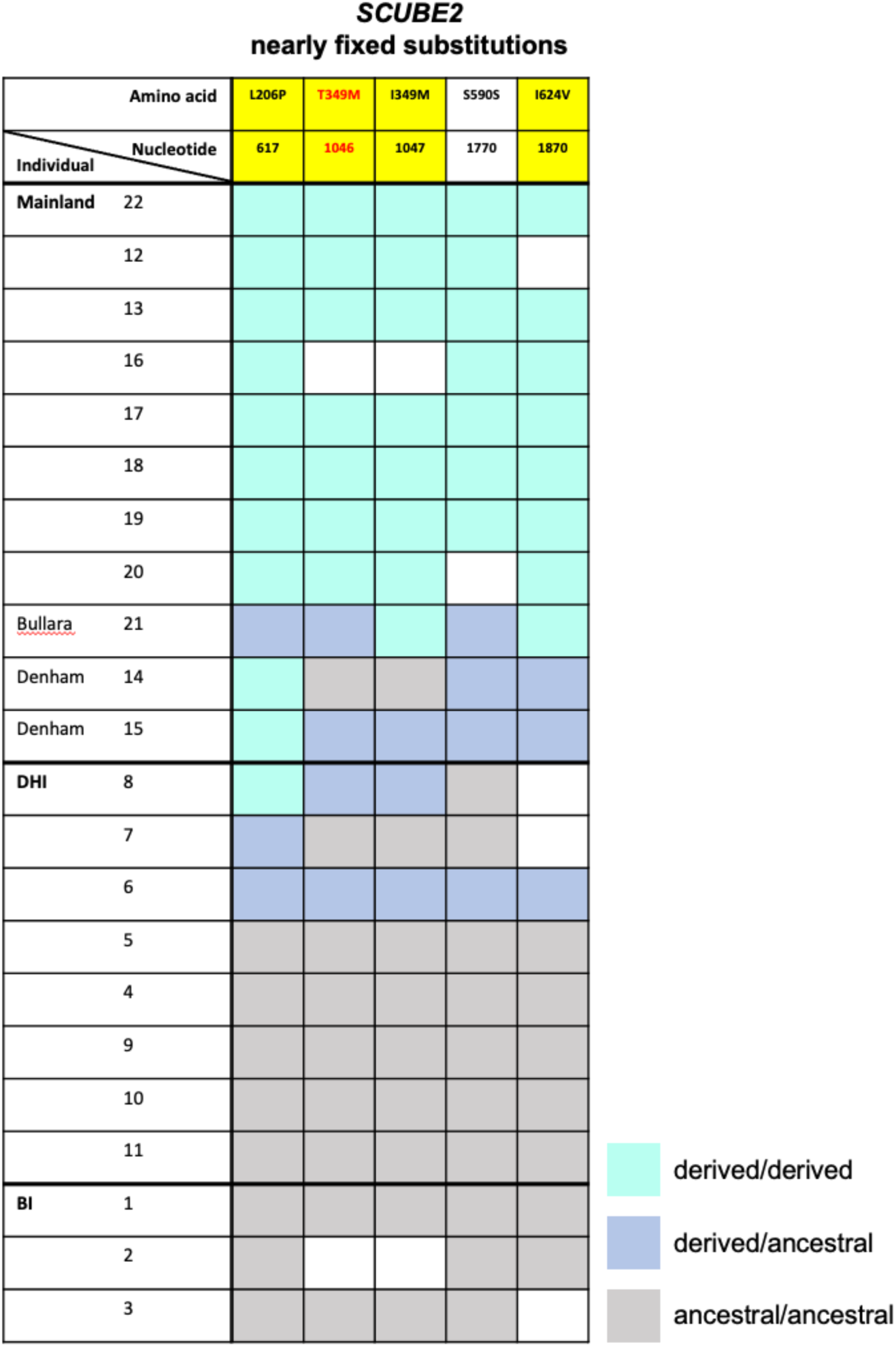
Genotypes at five nearly fixed nucleotide substitutions of the *SCUBE2* coding region between black (Dirk Hartog Island, DHI and Barrow Island, BI) and blue (mainland) *Malurus leucopterus* subspecies. The positions of nucleotide and amino acid substitutions are indicated at the top. Yellow indicates missense mutation. The red font indicates substitution of amino acid with different property. Each row represents one individual. Light blue, black, and dark blue indicate positions homozygous for the derived allele, homozygous for the ancestral allele, and heterozygous for both alleles, respectively. White denotes missing data.

## STAR METHODS

### Method details

#### Sample collection and DNA extraction

Samples from three subspecies of *Malurus leucopterus* were collected, including *M. leucopterus leuconotus* on mainland Australia (n = 11), *M. leucopterus leucopterus* on Dirk Hartog Island (n = 8), and *M. leucopterus edouardi* on Barrow Island (n = 3) (Fig. 1A; Supplementary Table S1). In total, 22 *M. leucopterus* males were included in this study, with 11 mainland individuals displayed a blue plumage and 11 insular individuals on island with black plumage. The two sister species *M. melanocephalus* (n = 11) and *M. alboscapulatus* (n = 11) were also sampled (Supplementary Table S1). One individual of *M. splendens*, a member of blue wrens that are sister to the bi-colored wrens, were included. *Malurus lamberti* and *Stipiturus malachurus* were also included as outgroups (Supplementary Table S1). Samples were collected from the field or requested from museum collections (Supplementary Table S1). We isolated genomic DNA using DNeasy Blood and Tissue Kit (Qiagen, Hilden, Germany) following the manufacturer’s protocol. We confirmed the sex of all individuals to be male using published primers targeting CHD1 genes (2550F & 2718R)^75^ for PCR and measured DNA concentration with a Qubit dsDNA HS Assay Kit (Invitrogen, Carlsbad, USA). Sample collection of *M. leucopterus* were approved by the Cornell University Institutional Animal Care and Use Committee (IACUC 2009-0105) and the James Cook University Animal Ethics Committee (A2100), and was performed with authorization of the Australian Bird and Bat Banding Scheme (#1965) and the Western Australia Department of Parks and Wildlife (BB003372). Sample collection of *M. alboscapulatus* was conducted under the approval of IACUC #0395 and ASAF #04573 and the Australian Bird and Bat Banding Scheme, and with annual permits from the Papua New Guinea Department of Environment and Conservation, permissions from the provincial governments of Milne Bay, Western, and Madang Provinces.

### Data generation and processing

#### Reference genome sequencing and assembly

The genome of one individual of each species was sequenced and assembled. We performed whole-genome library preparation, sequencing, and assembly following Sin et al.^76,77^ and Grayson et al.^78^. In brief, a DNA fragment library of 220 bp insert size was prepared for *M. leucopterus leuconotus* (from Queensland, Australia; Table S1), *M. splendens*, *M. lamberti*, and *S. malachurus* using the PrepX ILM 32i DNA Library Kit (Takara), and mate-pair libraries of 3 kb insert size were prepared for *M. leucopterus* and *Malurus splendens* using the Nextera Mate Pair Sample Preparation Kit (cat. No. FC-132-1001, Illumina). Mate-pair library of 6 kb insert size were also prepared for *M. leucopterus*. We performed DNA shearing for the fragment and mate-pair library preparations using Covaris S220. We used the 0.75% agarose cassette in the Pippin Prep (Sage Science) for size selection of the mate-pair library (target size 3 or 6 kb, “Tight” mode). We then assessed fragment and mate-pair library qualities using the High Sensitivity D1000 ScreenTape for the Tapestation (Agilent) and High Sensitivity DNA Kit for the Bioanalyzer (Agilent), respectively, and quantified the libraries with qPCR (KAPA library quantification kit) prior to sequencing. We sequenced the libraries on an Illumina HiSeq 2500 instrument (High Output 250 kit, PE 125 bp reads) at the Bauer Core facility at Harvard University to ∼65x coverage. We assessed the quality of the sequencing data using FastQC and removed adapters using Trimmomatic^79^. We performed *de novo* genome assembly using AllPaths-LG v52488^80^ for *M. leucopterus* and *M. splendens*. We estimated the completeness of the assembled genomes with BUSCO v2.0^81^ (Table S3). The *de novo* genomes of *M. alboscapulatus* (GCA_025434525.1)^82^ and *M. melanocephalus* (GCA_030028575.1)^83^ included in the analysis of this study were sequenced and assembled using the same strategies.

#### Genome annotation

We annotated the *M. leucopterus* genome following Sin et al.^84^. We used MAKER v2.31.8^85^, combining *ab initio* gene prediction with protein-based evidence from 16 other vertebrates (10 birds, 3 reptiles, 2 mammals, and 1 fish species) and gene predictions from *Gallus gallus*. The genome annotation using MAKER identified a total of 17,443 gene models. We functionally annotated the genome to identify putative gene function and protein domains using NCBI BLAST+ and the UniProt/Swiss-Prot set of proteins. We used BLASTP on the list of proteins identified by MAKER with an *e*-value of 1e-6.

#### Pseudochromosome reconstruction

We used Satsuma^86^ and MUMmer^87^ to align all assembled scaffolds in the *M. leucopterus* genome against the zebra finch (*Taeniopygia guttata*) genome (bTaeGut1_v1; GCF_003957565.1). A total of 312 sex-linked scaffolds – accounting for ∼9.08% of *Malurus leucopterus* draft genome – aligned to the Z chromosome. The aligned scaffolds of the *M. leucopterus* draft genome were reordered to form pseudochromosomes.

#### Whole-genome resequencing library preparation

To determine the genomic region(s) associated with male ornamentation polymorphism, we performed whole-genome resequencing at ∼4× depth-of-coverage for the 22 *M. leucopterus*. We also resequenced 11 samples of each of the two sister species, *M. melanocephalus* and *M. alboscapulatus*. DNA fragment libraries of 220 bp insert size were prepared using the protocol mentioned in the whole-genome library preparation section. All libraries were multiplexed in equimolar ratio and sequenced using an Illumina HiSeq instrument (High Output 250 kit, PE 125 bp reads). Preliminary quality assessment was performed using FastQC (www.bioinformatics.babraham.ac.uk/projects/fastqc).

### Data analysis

#### Whole-genome resequencing data preprocessing

We followed GATK best practices^88^ using Picard Tools v. 2.20.6 (broadinstitute.github.io/picard/) and Genome Analysis Toolkit (GATK) 3.8^89^ to preprocess the data. We removed adapters of the resequencing data using NGmerge v0.3^90^ and Picard Tools. We used BWA-MEM 0.7.13; ^91^ to map reads to the *M. leucopterus* genome assembly. The BAM files were merged, duplicates marked, sorted, and validated using Picard Tools. The reads were realigned around indels using GATK and undergone an initial variant calling using GATK HaplotypeCaller. We then applied a base quality score recalibration (BQSR) on SNPs and indels. The recalibrated BAM files were then passed to ANGSD v0.921^92^ to estimate genotype likelihood for final SNP calling, by first calculating the site allele frequency for each species/subspecies with the following settings: -skipTriAllelic 1 - remove_bads 1 -trim 0 -minMapQ 20 -minQ 20 -uniqueOnly 1 -baq 1 -C 50 - only_proper_pairs 0 -doCounts 1 -GL 1 -doMaf 1 -doPost 1 -doMajorMinor 1 -dobcf 1 -- ignore-RG 0 -dogeno 1 -doSaf 1, and the output was converted to VCF format using a custom R script (github.com/rcristofari/RAD-Scripts/blob/master/angsd2vcf.R). The VCF files from ANGSD were used for summary statistic calculations unless otherwise specified.

#### Inference of demographic history

To investigate the historical change in effective population size (*N*_e_) in *M. leucopterus* we used the Pairwise Sequential Markovian Coalescent (PSMC) model^93^ based on the diploid whole-genome sequence to reconstruct the population history. We filtered the raw sequencing data of the *M. leucopterus leuconotus* individual that we used for whole-genome assembly using fastp^94^ and removed sex-linked scaffolds. The settings for the PSMC atomic time intervals were “4+25*2+4+6”. We performed 100 bootstraps to compute the variance in estimate of *N*_e_. We used the estimated mutation rate of 3.3×10^-3^ substitutions per site per million years Passeriformes; ^95^. The estimated generation time is two years^96,97^.

#### Phylogenomic analysis and PCA

This VCF file was first formatted using a script vcf2phylip.py (https://github.com/edgardomortiz/vcf2phylip) for phylogenetic reconstruction using IQTREE 2.2.0^98^. Bi-allelic SNPs thinned by intervals of 5 kb, resulting in 168,378 polymorphic sites, were concatenated and used for Maximum Likelihood (ML) tree reconstruction with 1000 bootstraps. The best substitution model was selected using ModelFinder^99^. We also employed +ASC model to correct ascertainment bias as SNP data did not contain invariant sites, and we appended -bnni to the regular UFBoot command to reduce the risk of overestimating branch support^100^. Phylogeny was also inferred using the coalescent-based SNAPP package^101^ in BEAST2^102^ based on 6,790 genome-wide SNPs, with one SNP sampled from each 200 kb window. Two millions Monte Carlo Markov Chain (MCMC) generations were run with trees sampled every 1,000 generations. The resulting tree was visualized using DensiTree^103^. We also used ngsTools^104^ to generate a covariance matrix based on genotype-likelihoods for PCA.

#### Ancestral state inference

The ancestral genome of *M. leucopterus*, *M. melanocephalus*, and *M. alboscapulatus* was reconstructed using the software est-sfs v2.0395^105^ based on resequencing data, by setting these species in the bi-colored clade as ingroups and *M. splendens*, *M. lamberti*, and *S. malachurus* as outgroups, ordered from the most to the least closely related based on our inferred species trees. The software was designed to infer the unfolded SFS but it also output the allelic state probabilities of each site, which we further used to reconstruct the ancestral genome sequence. The inferred ancestral genome was then used in downstream genomic analyses of the resequenced samples to polarize alleles.

#### Summary statistic calculations and characterization of regions of divergence

We used the output from ANGSD and generated the site allele frequencies (SAF) (with *- doSaf*), which was then used to compute the unfolded SFS and to estimate various statistics. Alleles were polarized using the ancestral genome. We then obtained joint frequency spectrums for each between-subspecies comparison using realSFS, and then estimated *F*_ST_. We calculated *F*_ST_ for the blue and black *M. leucopterus* comparison. We calculated *F*_ST_ in overlapping sliding windows of 50 kb with 10 kb steps. Windows with *F*_ST_ above the 99.9^th^ percentile of autosomal values were considered regions of divergence and further examined for gene content.

Genetic diversity (π) and Fay and Wu’s H^106^ were also calculated for sliding windows across the genome using ANGSD. A significant negative value of Fay and Wu’s H (<99th percentile)^33^ indicates an excess of high-frequency derived SNPs due to selective sweep. We used SweepFinder2^107^ to detect signature of selective sweep in the regions of divergence within *M. leucopterus* subspecies with blue or black plumage. *Stipiturus malachurus* was used as the outgroup for this analysis, and the allele frequency within target group was calculated using VCFtools^108^. We estimated the composite likelihood ratio (CLR) statistics every 1000 bp. Elevated CLR values estimated by SweepFinder2 indicates a signature of selection sweep. We also conducted genome-wide association analysis (GWAS) based on allelic counts of each locus in Plink v1.90b6.26^109^ to identify the SNPs associated with the plumage color differentiation between black and blue *M. leucopterus*. Loci with a minor allele frequency lower than 0.05 and violated Hardy-Weinberg equilibrium (p ≤ 1e-6) were excluded for this analysis. To test if the candidate genomic regions underlying plumage color difference between blue and black *M. leucopterus* were introgressed from blue fairywren species that are sister to the bi-colored wrens, we calculated the relative node depth (RND)^110^ between blue and black *M. leucopterus*, using *M. alboscapulatus* as the outgroup. The RND was calculated as absolute nucleotide divergence (*D_XY_*) _(between black and blue *M. leucopterus*)_/([*D_XY_* _(between black *M. leucopterus* and *M. alboscapulatus*)_ + *D_XY_* _(between blue *M. leucopterus* and *M. alboscapulatus*)_]/2). *D_XY_* was calculated using pixy^111^ with VCF files generated by GATK’s GenotypeGVCFs, including variant and invariant sites. RND values are expected to lie between 0 and 1 for non-introgressed genomic regions, while for introgressed regions RND values should be well above 1^112^.

In the genomic regions of divergence, we examined all the annotated genes and identified those that are potentially relevant to plumage color development, such as genes related to pigment development or feather/keratinocyte development. For each highly divergent region with relevant candidate genes and a signature of selective sweep, we reconstructed the gene tree based on concatenated SNPs (2,355 and 666 SNPs for the *ASIP* and *SCUBE2* regions, respectively) using IQTREE 2.2.0^98^, with 1000 bootstraps. The best substitution model was selected using ModelFinder^99^. We also inferred the phylogeny using SNAPP^101^. Two millions Monte Carlo Markov Chain (MCMC) generations were run with trees sampled every 1,000 generations. Topology discordance between the species tree and trees of the regions with a selective sweep signature was tested using the Shimodaira-Hasegawa test (SH test)^113^ with 1,000 bootstrap replicates. We also visualized the genotypes using RectChr (github.com/BGI-shenzhen/RectChr). To perform gene enrichment analysis, we first extracted protein sequences from the genome annotation file and functionally annotated the sequences using eggNOG-mapper v2^114^. The genes in the top 99.9^th^ percentile *F*_ST_ peaks in the black and blue subspecies comparison were used to search for enriched pathways using clusterProfiler 4.0^115^.

To study the evolution of plumage color across *Malurus*, we investigated the SNP loci that had strong association with the plumage color difference between black and blue *M. leucopterus* in all studied species, including other bi-colored wrens and outgroup species. We first performed association analysis^109^ between the blue *M. leucopterus leuconotus* and black *M. leucopterus leucopterus* from Dirk Hartog Island, excluding those black individuals from the Barrow Island, and extracted those SNPs with strong association [i.e. −log(*P*)>6] with the color polymorphism. We then used RectChr to plot the genotype of these SNP loci in all individuals and species, including *M. leucopterus edouardi* from Barrow Island. We also visualized the allele frequencies of these loci in all *M. leucopterus* subspecies and other species.

We also examined the coding sequence (CDS) of the candidate genes for fixed or nearly fixed nucleotide substitutions. We used SnpEff^116^ to annotate the variants of the candidate genes and to differentiate synonymous and non-synonymous substitutions. Allele frequency at each site was calculated using VCFtools^108^.

### Resource availability

#### Lead contact

Further information and requests for resources and reagents should be directed to and will be fulfilled by the lead contact, Simon Yung Wa Sin.

#### Materials availability

This study did not generate new unique reagents.

#### Data and code accessibility

All genome assemblies and raw sequence read data will be deposited to the National Center for Biotechnology Information (NCBI). Accession number will be provided upon acceptance of the manuscript. All software are cited in the methods and are publicly available.

## References

1. Baeckens, S., and Van Damme, R. (2020). The island syndrome. Curr. Biol. 30, R338–R339.

2. Lamichhaney, S., Han, F., Berglund, J., Wang, C., Almén, M.S., Webster, M.T., Grant, B.R., Grant, P.R., and Andersson, L. (2016). A beak size locus in Darwin’s finches facilitated character displacement during a drought. Science 352, 470–474.

3. Cerca, J., Cotoras, D.D., Bieker, V.C., De-Kayne, R., Vargas, P., Fernández-Mazuecos, M., López-Delgado, J., White, O., Stervander, M., and Geneva, A.J. (2023). Evolutionary genomics of oceanic island radiations. Trends Ecol. Evol.

4. Uy, J.A.C., Cooper, E.A., Cutie, S., Concannon, M.R., Poelstra, J.W., Moyle, R.G., and Filardi, C.E. (2016). Mutations in different pigmentation genes are associated with parallel melanism in island flycatchers. Proc. R. Soc. Lond. B Biol. Sci. 283, 20160731.

5. Romano, A., Séchaud, R., and Roulin, A. (2021). Evolution of wing length and melanin-based coloration in insular populations of a cosmopolitan raptor. J. Biogeogr. 48, 961–973.

6. Campagna, L., Mo, Z., Siepel, A., and Uy, J.A.C. (2022). Selective sweeps on different pigmentation genes mediate convergent evolution of island melanism in two incipient bird species. Plos Genetics 18, e1010474.

7. Driskell, A.C., Pruett-Jones, S., Tarvin, K.A., and Hagevik, S. (2002). Evolutionary relationships among blue-and black-plumaged populations of the white-winged fairy-wren (Malurus leucopterus). Aust. J. Zool. 50, 581–595.

8. Uy, J.A.C., Cooper, E.A., and Chaves, J.A. (2019). Convergent melanism in populations of a Solomon Island flycatcher is mediated by unique genetic mechanisms. Emu-Austral Ornithology 119, 242–250.

9. Buades, J.M., Rodríguez, V., Terrasa, B., Perez-Mellado, V., Brown, R.P., Castro, J.A., Picornell, A., and Ramon, M. (2013). Variability of the mc1r Gene in Melanic and Non-Melanic Podarcis lilfordi and Podarcis pityusensis from the Balearic Archipelago. PloS one 8, e53088.

10. Castilla, A. (1994). A case of melanism in a population of the insular lizard Podarcis hispanica atrata. Bolletí de la Societat d’Història Natural de les Balears, 175–179.

11. Ritchie, J. (1978). Melanism in Oedaleus senegalensis and other oedipodines (Orthoptera, Acrididae). J. Nat. Hist. 12, 153–162.

12. Roulin, A. (2014). Melanin-based colour polymorphism responding to climate change. Global Change Biology 20, 3344–3350.

13. Goldstein, G., Flory, K.R., Browne, B.A., Majid, S., Ichida, J.M., and Burtt Jr, E.H. (2004). Bacterial degradation of black and white feathers. The Auk 121, 656–659.

14. Bonser, R.H. (1995). Melanin and the abrasion resistance of feathers. The Condor 97, 590–591.

15. Margalida, A., Negro, J.J., and Galván, I. (2008). Melanin-based color variation in the bearded vulture suggests a thermoregulatory function. Comparative Biochemistry and Physiology Part A: Molecular & Integrative Physiology 149, 87–91.

16. Rathburn, M.K., and Montgomerie, R. (2003). Breeding biology and social structure of White-winged Fairy-wrens (Malurus leucopterus): comparison between island and mainland subspecies having different plumage phenotypes. Emu 103, 295–306.

17. Surmacki, A., Minias, P., and Kudelska, K. (2021). Occurrence and function of melanin-based grey coloration in Western Palaearctic songbirds (Aves: Passeriformes). Ibis 163, 390–406.

18. Doucet, S., Shawkey, M., Rathburn, M., Mays Jr, H., and Montgomerie, R. (2004). Concordant evolution of plumage colour, feather microstructure and a melanocortin receptor gene between mainland and island populations of a fairy–wren. Proceedings of the Royal Society of London. Series B: Biological Sciences 271, 1663–1670.

19. Schodde, R. (1982). The Fairy-wrens: a monograph of the Maluridae. Lansdowne, Melbourne.

20. Ford, J. (1987). Minor isolates and minor geographical barriers in avian speciation in continental Australia. Emu 87, 90–102.

21. Mundy, N.I. (2005). A window on the genetics of evolution: MC1R and plumage colouration in birds. Proc. R. Soc. Lond. B Biol. Sci. 272, 1633–1640.

22. Toomey, M.B., Marques, C.I., Araújo, P.M., Huang, D., Zhong, S., Liu, Y., Schreiner, G.D., Myers, C.A., Pereira, P., and Afonso, S. (2022). A mechanism for red coloration in vertebrates. Curr. Biol. 32, 4201–4214. e4212.

23. Cooke, T.F., Fischer, C.R., Wu, P., Jiang, T.-X., Xie, K.T., Kuo, J., Doctorov, E., Zehnder, A., Khosla, C., and Chuong, C.-M. (2017). Genetic mapping and biochemical basis of yellow feather pigmentation in budgerigars. Cell 171, 427–439. e421.

24. Price-Waldman, R., and Stoddard, M.C. (2021). Avian coloration genetics: recent advances and emerging questions. J. Hered. 112, 395–416.

25. Prum, R.O., and Torres, R.H. (2003). A Fourier tool for the analysis of coherent light scattering by bio-optical nanostructures. Integrative and Comparative Biology 43, 591–602.

26. Shawkey, M.D., and D’Alba, L. (2017). Interactions between colour-producing mechanisms and their effects on the integumentary colour palette. Philosophical Transactions of the Royal Society B: Biological Sciences 372, 20160536.

27. Saranathan, V., and Finet, C. (2021). Cellular and developmental basis of avian structural coloration. Curr. Opin. Genet. Dev. 69, 56–64.

28. Prum, R.O., Dufresne, E.R., Quinn, T., and Waters, K. (2009). Development of colour-producing β-keratin nanostructures in avian feather barbs. J. Royal Soc. Interface 6, S253–S265.

29. Fan, M., D’alba, L., Shawkey, M.D., Peters, A., and Delhey, K. (2019). Multiple components of feather microstructure contribute to structural plumage colour diversity in fairy-wrens. Biol. J. Linn. Soc. 128, 550–568.

30. Driskell, A.C., Prum, R.O., and Pruett-Jones, S. (2010). The evolution of black plumage from blue in Australian fairy-wrens (Maluridae): genetic and structural evidence. J. Avian Biol. 41, 505–514.

31. Walsh, J., Campagna, L., Feeney, W.E., King, J., and Webster, M.S. (2021). Patterns of genetic divergence and demographic history shed light on island-mainland population dynamics and melanic plumage evolution in the white-winged Fairywren. Evolution 75, 1348–1360.

32. Robert, A., Peona, V., and Ottenburghs, J. (2021). Digest: Population genomics reveals convergence toward melanism in different island populations. Evolution 75, 1582–1584.

33. Sterken, R., Kiekens, R., Coppens, E., Vercauteren, I., Zabeau, M., Inze, D., Flowers, J., and Vuylsteke, M. (2009). A population genomics study of the Arabidopsis core cell cycle genes shows the signature of natural selection. The Plant Cell 21, 2987–2998.

34. Gantz, I., and Fong, T.M. (2003). The melanocortin system. American Journal of Physiology-Endocrinology and Metabolism.

35. Lin, S., Foley, J., Jiang, T., Yeh, C., Wu, P., Foley, A., Yen, C., Huang, Y., Cheng, H., and Chen, C. (2013). Topology of feather melanocyte progenitor niche allows complex pigment patterns to emerge. Science 340, 1442–1445.

36. D’Alba, L., and Shawkey, M.D. (2019). Melanosomes: biogenesis, properties, and evolution of an ancient organelle. Physiol. Rev. 99, 1–19.

37. Nadeau, N.J., Minvielle, F., Ito, S.i., Inoue-Murayama, M., Gourichon, D., Follett, S.A., Burke, T., and Mundy, N.I. (2008). Characterization of Japanese quail yellow as a genomic deletion upstream of the avian homolog of the mammalian ASIP (agouti) gene. Genetics 178, 777–786.

38. Norris, B.J., and Whan, V.A. (2008). A gene duplication affecting expression of the ovine ASIP gene is responsible for white and black sheep. Genome Res. 18, 1282–1293.

39. Kratochwil, C.F. (2019). Molecular mechanisms of convergent color pattern evolution. Zoology 134, 66–68.

40. Robic, A., Morisson, M., Leroux, S., Gourichon, D., Vignal, A., Thebault, N., Fillon, V., Minvielle, F., Bed’Hom, B., and Zerjal, T. (2019). Two new structural mutations in the 5′ region of the ASIP gene cause diluted feather color phenotypes in Japanese quail. Genetics Selection Evolution 51, 1–10.

41. Wang, S., Rohwer, S., de Zwaan, D.R., Toews, D.P., Lovette, I.J., Mackenzie, J., and Irwin, D. (2020). Selection on a small genomic region underpins differentiation in multiple color traits between two warbler species. Evolution Letters 4, 502–515.

42. Goutte, S., Hariyani, I., Utzinger, K.D., Bourgeois, Y., and Boissinot, S. (2022). Genomic Analyses Reveal Association of ASIP with a Recurrently evolving Adaptive Color Pattern in Frogs. Mol. Biol. Evol. 39, msac235.

43. Semenov, G.A., Linck, E., Enbody, E.D., Harris, R.B., Khaydarov, D.R., Alström, P., Andersson, L., and Taylor, S.A. (2021). Asymmetric introgression reveals the genetic architecture of a plumage trait. Nature Communications 12, 1019.

44. Jakobs, P., Schulz, P., Schürmann, S., Niland, S., Exner, S., Rebollido-Rios, R., Manikowski, D., Hoffmann, D., Seidler, D.G., and Grobe, K. (2017). Ca2+ coordination controls sonic hedgehog structure and its Scube2-regulated release. J. Cell Sci. 130, 3261–3271.

45. Tsai, M.-T., Cheng, C.-J., Lin, Y.-C., Chen, C.-C., Wu, A.-R., Wu, M.-T., Hsu, C.-C., and Yang, R.-B. (2009). Isolation and characterization of a secreted, cell-surface glycoprotein SCUBE2 from humans. Biochem. J. 422, 119–128.

46. Cooper, R.L., and Milinkovitch, M.C. (2023). Transient agonism of the sonic hedgehog pathway triggers a permanent transition of skin appendage fate in the chicken embryo. Science Advances 9, eadg9619.

47. Ting-Berreth, S.A., and Chuong, C.M. (1996). Sonic Hedgehog in feather morphogenesis: induction of mesenchymal condensation and association with cell death. Dev. Dyn. 207, 157–170.

48. Jung, H.-S., Francis-West, P.H., Widelitz, R.B., Jiang, T.-X., Ting-Berreth, S., Tickle, C., Wolpert, L., and Chuong, C.-M. (1998). Local inhibitory action of BMPs and their relationships with activators in feather formation: implications for periodic patterning. Dev. Biol. 196, 11–23.

49. Chuong, C.-M., Chodankar, R., Widelitz, R.B., and Jiang, T.-X. (2000). Evo-devo of feathers and scales: building complex epithelial appendages. Curr. Opin. Genet. Dev. 10, 449.

50. Yu, M., Wu, P., Widelitz, R.B., and Chuong, C.-M. (2002). The morphogenesis of feathers. Nature 420, 308–312.

51. Tanaka, Y., Nakanishi, H., Kakunaga, S., Okabe, N., Kawakatsu, T., Shimizu, K., and Takai, Y. (2003). Role of nectin in formation of E-cadherin–based adherens junctions in keratinocytes: analysis with the N-cadherin dominant negative mutant. Mol. Biol. Cell 14, 1597–1609.

52. Yoshida, T., Takai, Y., and Thesleff, I. (2014). Cooperation of Nectin-1 and Nectin-3 is required for maintenance of epidermal stratification and proper hair shaft formation in the mouse. Developmental Biology Journal 2014.

53. Adams, J.C., and Lawler, J. (2004). The thrombospondins. The international journal of biochemistry & cell biology 36, 961–968.

54. Adams, J.C., and Lawler, J. (2011). The thrombospondins. Cold Spring Harbor perspectives in biology 3, a009712.

55. Mäemets-Allas, K., Klaas, M., Cárdenas-León, C.G., Arak, T., Kankuri, E., and Jaks, V. (2023). Stimulation with THBS4 activates pathways that regulate proliferation, migration and inflammation in primary human keratinocytes. Biochem. Biophys. Res. Commun. 642, 97–106.

56. Haines, P. (2014). Hybrid fairywren at Hart Lagoon, River Murray, South Australia. S. Aust. Ornithol. 39, 87–89.

57. Ross, M., and Briggs, A. (2022). Behaviour of probable hybrid Red-backed x Superb Fairy-wrens at Gladstone, Queensland. Australian Field Ornithology 39, 76–81.

58. Welklin, J.F., Johnson, A.E., Black, A., Nye, G., Sramek, P., Michalek, B., Ross, M., Newport, A., Walker, D., and Welburn, T. (2022). New records of hybridisation in Australian Fairy-wrens’ Malurus’ spp. Australian Field Ornithology 39, 63–75.

59. Shawkey, M.D., Estes, A.M., Siefferman, L., and Hill, G.E. (2005). The anatomical basis of sexual dichromatism in non-iridescent ultraviolet-blue structural coloration of feathers. Biol. J. Linn. Soc. 84, 259–271.

60. Karubian, J. (2002). Costs and benefits of variable breeding plumage in the red- backed fairy-wren. Evolution 56, 1673–1682.

61. Colombelli-Négrel, D., Schlotfeldt, B.E., and Kleindorfer, S. (2009). High levels of extra-pair paternity in Superb Fairy-wrens in South Australia despite low frequency of auxiliary males. Emu-Austral Ornithology 109, 300–304.

62. Cockburn, A., Brouwer, L., Double, M.C., Margraf, N., and van de Pol, M. (2013). Evolutionary origins and persistence of infidelity in Malurus: the least faithful birds. Emu 113, 208–217.

63. Lindsay, W.R., Webster, M.S., and Schwabl, H. (2011). Sexually selected male plumage color is testosterone dependent in a tropical passerine bird, the red-backed fairy-wren (Malurus melanocephalus). PloS one 6, e26067.

64. Friedman, N., and Remeš, V. (2015). Rapid evolution of elaborate male coloration is driven by visual system in A ustralian fairy-wrens (M aluridae). J. Evol. Biol. 28, 2125–2135.

65. Webster, M.S., Varian, C.W., and Karubian, J. (2008). Plumage color and reproduction in the red-backed fairy-wren: why be a dull breeder? Behav. Ecol. 19, 517–524.

66. Montgomerie, R. (2006). Cosmetic and adventitious colors. Bird coloration 1, 399–427.

67. Karubian, J., and Alvarado, A. (2003). Testing the function of petal-carrying in the Red-backed Fairy-wren (Malurus melanocephalus). Emu 103, 87–92.

68. Hejase, H.A., Salman-Minkov, A., Campagna, L., Hubisz, M.J., Lovette, I.J., Gronau, I., and Siepel, A. (2020). Genomic islands of differentiation in a rapid avian radiation have been driven by recent selective sweeps. Proceedings of the National Academy of Sciences 117, 30554–30565.

69. Endler, J.A. (1990). On the measurement and classification of colour in studies of animal colour patterns. Biol. J. Linn. Soc. 41, 315–352.

70. Omland, K.E. (1997). Examining two standard assumptions of ancestral reconstructions: repeated loss of dichromatism in dabbling ducks (Anatini). Evolution 51, 1636–1646.

71. Frankham, R. (1997). Do island populations have less genetic variation than mainland populations? Heredity 78, 311–327.

72. Griffith, S.C. (2000). High fidelity on islands: a comparative study of extrapair paternity in passerine birds. Behav. Ecol. 11, 265–273.

73. Maia, R., Rubenstein, D.R., and Shawkey, M.D. (2013). Key ornamental innovations facilitate diversification in an avian radiation. Proceedings of the National Academy of Sciences 110, 10687–10692.

74. Collins, P. (1995). Variation in the plumage of the White-winged Fairy-wren’Malurus leucopterus’. Australian Bird Watcher 16, 130–131.

75. Fridolfsson, A.K., and Ellegren, H. (1999). A simple and universal method for molecular sexing of non-ratite birds. J. Avian Biol. 30, 116–121.

76. Sin, S.Y.W., Lu, L., and Edwards, S.V. (2020). De Novo assembly of the Northern Cardinal (*Cardinalis cardinalis*) genome reveals candidate regulatory regions for sexually dichromatic red plumage coloration. G3: Genes, Genomes, Genetics 10, 3541–3548.

77. Huynh, S., Cloutier, A., Chen, G., Chan, D.T.C., Lam, D.K., Huyvaert, K.P., Sato, F., Edwards, S.V., and Sin, S.Y.W. (2023). Whole-genome analyses reveal past population fluctuations and low genetic diversities of the North Pacific albatrosses. Mol. Biol. Evol. 40, msad155.

78. Grayson, P., Sin, S.Y.W., Sackton, T.B., and Edwards, S.V. (2017). Comparative genomics as a foundation for evo-devo studies in birds (Humana Press, New York, NY).

79. Bolger, A., Lohse, M., and Usadel, B. (2014). Trimmomatic: a flexible trimmer for Illumina sequence data. Bioinformatics 30, 2114–2120.

80. Gnerre, S., MacCallum, I., Przybylski, D., Ribeiro, F., Burton, J., Walker, B., Sharpe, T., Hall, G., Shea, T., Sykes, S., and Berlin, A. (2011). High-quality draft assemblies of mammalian genomes from massively parallel sequence data. Proceedings of the National Academy of Sciences 108, 1513–1518.

81. Simão, F.A., Waterhouse, R.M., Ioannidis, P., Kriventseva, E.V., and Zdobnov, E.M. (2015). BUSCO: assessing genome assembly and annotation completeness with single-copy orthologs. Bioinformatics 31, 3210–3212.

82. Enbody, E.D., Sin, S.Y., Boersma, J., Edwards, S.V., Ketaloya, S., Schwabl, H., Webster, M.S., and Karubian, J. (2022). The evolutionary history and mechanistic basis of female ornamentation in a tropical songbird. Evolution 76, 1720–1736.

83. Khalil, S., Enbody, E.D., Frankl-Vilches, C., Welklin, J.F., Koch, R.E., Toomey, M.B., Sin, S.Y.W., Edwards, S.V., Gahr, M., and Schwabl, H. (2023). Testosterone coordinates gene expression across different tissues to produce carotenoid-based red ornamentation. Mol. Biol. Evol., msad056.

84. Sin, S.Y.W., Cloutier, A., Nevitt, G., and Edwards, S.V. (2022). Olfactory receptor subgenome and expression in a highly olfactory procellariiform seabird. Genetics 220, iyab210.

85. Holt, C., and Yandell, M. (2011). MAKER2: an annotation pipeline and genome-database management tool for second-generation genome projects. BMC Bioinformatics 12, 491.

86. Grabherr, M.G., Russell, P., Meyer, M., Mauceli, E., Alföldi, J., Palma, F.D., and Lindblad-Toh, K. (2010). Genome-wide synteny through highly sensitive sequence alignment: Satsuma. Bioinformatics 26, 1145–1151.

87. Kurtz, S., Phillippy, A., Delcher, A.L., Smoot, M., Shumway, M., Antonescu, C., and Salzberg, S.L. (2004). Versatile and open software for comparing large genomes. Genome Biology 5, R12.

88. Van der Auwera, G.A., Carneiro, M.O., Hartl, C., Poplin, R., Del Angel, G., Levy- Moonshine, A., Jordan, T., Shakir, K., Roazen, D., and Thibault, J. (2013). From FastQ data to high-confidence variant calls: the genome analysis toolkit best practices pipeline. Current protocols in bioinformatics 43, 11.10. 11–11.10. 33.

89. McKenna, A., Hanna, M., Banks, E., Sivachenko, A., Cibulskis, K., Kernytsky, A., Garimella, K., Altshuler, D., Gabriel, S., Daly, M., and DePristo, M. (2010). The Genome Analysis Toolkit: a MapReduce framework for analyzing next-generation DNA sequencing data. Genome Res. 20, 1297–1303.

90. Gaspar, J.M. (2018). NGmerge: merging paired-end reads via novel empirically-derived models of sequencing errors. BMC Bioinformatics 19, 1–9.

91. Li, H., and Durbin, R. (2009). Fast and accurate short read alignment with Burrows– Wheeler transform. bioinformatics 25, 1754–1760.

92. Korneliussen, T., Albrechtsen, A., and Nielsen, R. (2014). ANGSD: analysis of next generation sequencing data. BMC Bioinformatics 15, 356.

93. Li, H., and Durbin, R. (2011). Inference of human population history from individual whole-genome sequences. Nature 475, 493–496.

94. Chen, S., Zhou, Y., Chen, Y., and Gu, J. (2018). fastp: an ultra-fast all-in-one FASTQ preprocessor. Bioinformatics 34, i884–i890.

95. Zhang, G., Li, C., Li, Q., Li, B., Larkin, D., Lee, C., Storz, J., Antunes, A., Greenwold, M., Meredith, R., and Ödeen, A. (2014). Comparative genomics reveals insights into avian genome evolution and adaptation. Science 346, 1311–1320.

96. Rowley, I., and Russell, E. (1995). The breeding biology of the White-winged Fairy-wren Malurus leucopterus leuconotus in a Western Australian coastal heathland. Emu-Austral Ornithology 95, 175–184.

97. Garnett, S.T. (2021). The action plan for Australian birds 2020 (CSIRO publishing).

98. Minh, B.Q., Schmidt, H.A., Chernomor, O., Schrempf, D., Woodhams, M.D., Von Haeseler, A., and Lanfear, R. (2020). IQ-TREE 2: new models and efficient methods for phylogenetic inference in the genomic era. Mol. Biol. Evol. 37, 1530–1534.

99. Kalyaanamoorthy, S., Minh, B.Q., Wong, T.K., Von Haeseler, A., and Jermiin, L.S. (2017). ModelFinder: fast model selection for accurate phylogenetic estimates. Nat. Meth. 14, 587–589.

100. Hoang, D.T., Chernomor, O., Von Haeseler, A., Minh, B.Q., and Vinh, L.S. (2018). UFBoot2: improving the ultrafast bootstrap approximation. Mol. Biol. Evol. 35, 518–522.

101. Bryant, D., Bouckaert, R., Felsenstein, J., Rosenberg, N.A., and RoyChoudhury, A. (2012). Inferring species trees directly from biallelic genetic markers: bypassing gene trees in a full coalescent analysis. Mol. Biol. Evol. 29, 1917–1932.

102. Bouckaert, R., Vaughan, T.G., Barido-Sottani, J., Duchêne, S., Fourment, M., Gavryushkina, A., Heled, J., Jones, G., Kühnert, D., and De Maio, N. (2019). BEAST 2.5: An advanced software platform for Bayesian evolutionary analysis. PLoS computational biology 15, e1006650.

103. Bouckaert, R.R. (2010). DensiTree: making sense of sets of phylogenetic trees. Bioinformatics 26, 1372–1373.

104. Fumagalli, M., Vieira, F.G., Linderoth, T., and Nielsen, R. (2014). ngsTools: methods for population genetics analyses from next-generation sequencing data. Bioinformatics 30, 1486–1487.

105. Keightley, P.D., and Jackson, B.C. (2018). Inferring the probability of the derived vs. the ancestral allelic state at a polymorphic site. Genetics 209, 897–906.

106. Fay, J.C., and Wu, C.-I. (2000). Hitchhiking under positive Darwinian selection. Genetics 155, 1405–1413.

107. DeGiorgio, M., Huber, C.D., Hubisz, M.J., Hellmann, I., and Nielsen, R. (2016). SweepFinder2: increased sensitivity, robustness and flexibility. Bioinformatics 32, 1895–1897.

108. Danecek, P., Auton, A., Abecasis, G., Albers, C., Banks, E., DePristo, M., Handsaker, R., Lunter, G., Marth, G., Sherry, S., and McVean, G. (2011). The variant call format and VCFtools. Bioinformatics 27, 2156–2158.

109. Purcell, S., Neale, B., Todd-Brown, K., Thomas, L., Ferreira, M., Bender, D., Maller, J., Sklar, P., Bakker, P.D., Daly, M., and Sham, P. (2007). PLINK: a tool set for whole-genome association and population-based linkage analyses. The American journal of human genetics 81, 559–575.

110. Feder, J.L., Xie, X., Rull, J., Velez, S., Forbes, A., Leung, B., Dambroski, H., Filchak, K.E., and Aluja, M. (2005). Mayr, Dobzhansky, and Bush and the complexities of sympatric speciation in Rhagoletis. Proceedings of the National Academy of Sciences 102, 6573–6580.

111. Korunes, K.L., and Samuk, K. (2021). pixy: Unbiased estimation of nucleotide diversity and divergence in the presence of missing data. Molecular ecology resources 21, 1359–1368.

112. Lopes, R.J., Johnson, J.D., Toomey, M.B., Ferreira, M.S., Araujo, P.M., Melo-Ferreira, J., Andersson, L., Hill, G.E., Corbo, J.C., and Carneiro, M. (2016). Genetic basis for red coloration in birds. Curr. Biol. 26, 1427–1434.

113. Shimodaira, H., and Hasegawa, M. (1999). Multiple comparisons of log-likelihoods with applications to phylogenetic inference. Mol. Biol. Evol. 16, 1114.

114. Cantalapiedra, C.P., Hernández-Plaza, A., Letunic, I., Bork, P., and Huerta-Cepas, J. (2021). eggNOG-mapper v2: functional annotation, orthology assignments, and domain prediction at the metagenomic scale. Mol. Biol. Evol. 38, 5825–5829.

115. Wu, T., Hu, E., Xu, S., Chen, M., Guo, P., Dai, Z., Feng, T., Zhou, L., Tang, W., and Zhan, L. (2021). clusterProfiler 4.0: A universal enrichment tool for interpreting omics data. The Innovation 2, 100141.

116. Cingolani, P., Platts, A., Wang, L., Coon, M., Nguyen, T., Wang, L., Land, S., Lu, X., and Ruden, D. (2012). A program for annotating and predicting the effects of single nucleotide polymorphisms, SnpEff: SNPs in the genome of Drosophila melanogaster strain w1118; iso-2; iso-3. Fly 6, 80–92.

